# Gut bacteria-derived succinate drives enteric neuron recovery following microbiota perturbation

**DOI:** 10.1101/2024.10.15.618589

**Authors:** Begüm Aydin, Izabela Mamede, Matthew R. Paul, Douglas Barrows, Joana Cardoso, Julia Deere, Huixian Li, Shanshan Qiao, Yelina Alvarez, Dong-Yoon Kim, Sofia Archontidi, Ved P. Sharma, Liwen Deng, Georgia Gunner, Marissa A. Scavuzzo, Gregory P. Donaldson, Isaac M. Chiu, Vassilis Pachnis, Chun-Jun Guo, Daniel Mucida

## Abstract

Enteric neurons control gut physiology by regulating peristalsis, nutrient absorption, and secretion^1^. Disruptions in microbial communities caused by antibiotics or enteric infections lead to loss of enteric neurons and long-term motility disorders^2–5^. However, little is known about the signals and underlying mechanisms of microbiota– neuron communication. We found that both enteric neurons and glia are lost after antibiotic treatment but recover when the pre-treatment microbiota is restored. Using murine gnotobiotic models, metabolomics, and murine and bacterial genetic approaches, we identified neurogenic bacterial species and their metabolite, succinate, as a microbiota-derived signal that induces enteric neuronal recovery in a tuft cell-dependent manner. Single-nucleus expression profiling during enteric neuron recovery revealed a precursor-like state marked by the expression of the neuronal gene, *Nav2*. Glial chromatin profiling and genetic fate mapping showed that Plp1+ glial cells acquire a Nav2-associated recovery state following microbial perturbation and contribute to neuronal recovery. In contrast, succinate recapitulated aspects of developmental neuron subtype diversification in the adult ENS and increased Nav2+ neurons independent of the Plp1 lineage, indicating that distinct microbe-dependent mechanisms converge on recovery-associated neuronal states. Our findings reveal a specific microbe-derived metabolite and the downstream cellular mechanisms involved in neuronal regeneration, with potential therapeutic implications for peripheral neuropathies.

## Background

The gastrointestinal (GI) tract is the largest barrier surface through which the body interacts with its external environment. This extensive interface hosts a vast array of immune cells, gut microbiota, and the enteric nervous system (ENS), a complex network of neurons and glial cells responsible for regulating essential gastrointestinal functions and responding to diverse environmental signals^1,6^. The intricate crosstalk between the ENS, gut immune system, and microbiota creates a dynamic microenvironment. However, the signals and mechanisms that mediate this communication remain poorly understood. Disruptions in this balance, often involving alterations in microbial communities, have been implicated in the pathogenesis of post-infectious neuropathies and neurodegenerative and neurodevelopmental disorders^7,8^. Most enteric neurons and glia are derived from neural crest progenitors that invade the gut during embryogenesis^9^, undergoing a finely tuned process of self-renewal and differentiation, and giving rise to diverse neuronal and glial cell types along the length of the gut^10^. A secondary wave of neurogenesis occurs postnatally^11^, coinciding with gut colonization by microbiota, which regulates the postnatal migration of enteric glial cells along the serosa-lumen axis^12^. Mice raised under germ-free (GF) conditions have a reduced number of enteric neurons in the ileum and colon, the regions where the density and diversity of the microbiota are highest in conventional mice^3^. This temporal and spatial association suggests a possible role for microbial signals in influencing ENS development and maintenance following microbial insults. While enteric neurogenesis is thought to cease by weaning under physiological conditions^11^, studies have demonstrated the remarkable plasticity of enteric glial cells, which can differentiate into neurons *in vitro* and *in vivo* following chemical injury or colitis^13–15^. Nonetheless, the molecular signals, particularly those derived from the microbiota, that govern ENS plasticity and the role of microbiota in ENS maintenance and regeneration remain largely unexplored. Here, we investigated the effects of microbiota on the ENS following microbial dysbiosis and colonization in GF mice to uncover the bacterial signals and cellular mechanisms that maintain postnatal ENS plasticity.

### Microbial manipulations mediate loss and recovery of the enteric nervous system

We have previously shown that microbial dysbiosis induced by antibiotic (Abx) treatment or *Salmonella enterica ΔspiB (Salmonella ΔspiB)* infection results in the loss of enteric neurons mediated by the caspase-11/NLRP6-dependent non-canonical inflammasome pathway^2,3^. Following acute treatment with a single Abx, ampicillin, in wild-type C57Bl/6 mice, we observed the loss of both enteric neurons and glia in the distal small intestine (**Fig. 1a, b**). While we observed a decrease in overall Sox10+ glia (**Fig. 1a**), we quantified only intraganglionic glial cells to avoid contributions from extraganglionic glia, whose numbers can fluctuate with imaged tissue thickness. The neuronal and glial loss observed with Abx treatment was also replicated in outbred Swiss Webster mice, indicating that it is not restricted to a specific inbred strain (**Fig. 1c**). These observations confirm ENS cell loss following Abx treatment^3,12,16,17^. To test whether both neurons and glia recover upon microbiota restoration, we performed fecal microbiota transfer (FMT) from specific pathogen-free (SPF) mice, housed either in-house or at Jackson Laboratory, to test recovery across distinct microbiota (**Fig. 1d, e, g**). Mice were treated with ampicillin via oral gavage for five days, and age-matched mice from the same donor genotype were used as fecal donors to minimize microbiota-related differences in all fecal transfer experiments. Ampicillin treatment led to dysbiosis, characterized by reduced overall diversity and increased *Bacteroides* abundance, as determined by 16S rRNA sequencing of fecal samples. (**Extended Data Fig. 1a**). Both neuron and glial cell numbers were recovered seven days after fecal transfer in both in-house and Jackson mice, reaching the levels observed at their steady state, as indicated by the gray-shaded area on individual graphs (**Fig. 1d–f**). The recovery in neuronal numbers also correlated with the recovery in bacterial diversity in steady-state (SPF) animals (**Extended Data Fig. 1a**). In addition, SPF FMT rescued transit time defects observed in Abx-treated mice (**Fig. 1g**). However, dysbiotic FMT taken from mice at the time of treatment did not rescue neuronal and glial numbers, unlike SPF FMT (**Fig. 1f**), corroborating the restoration of dysbiosis driving the observed recovery.

**Fig. 1.**
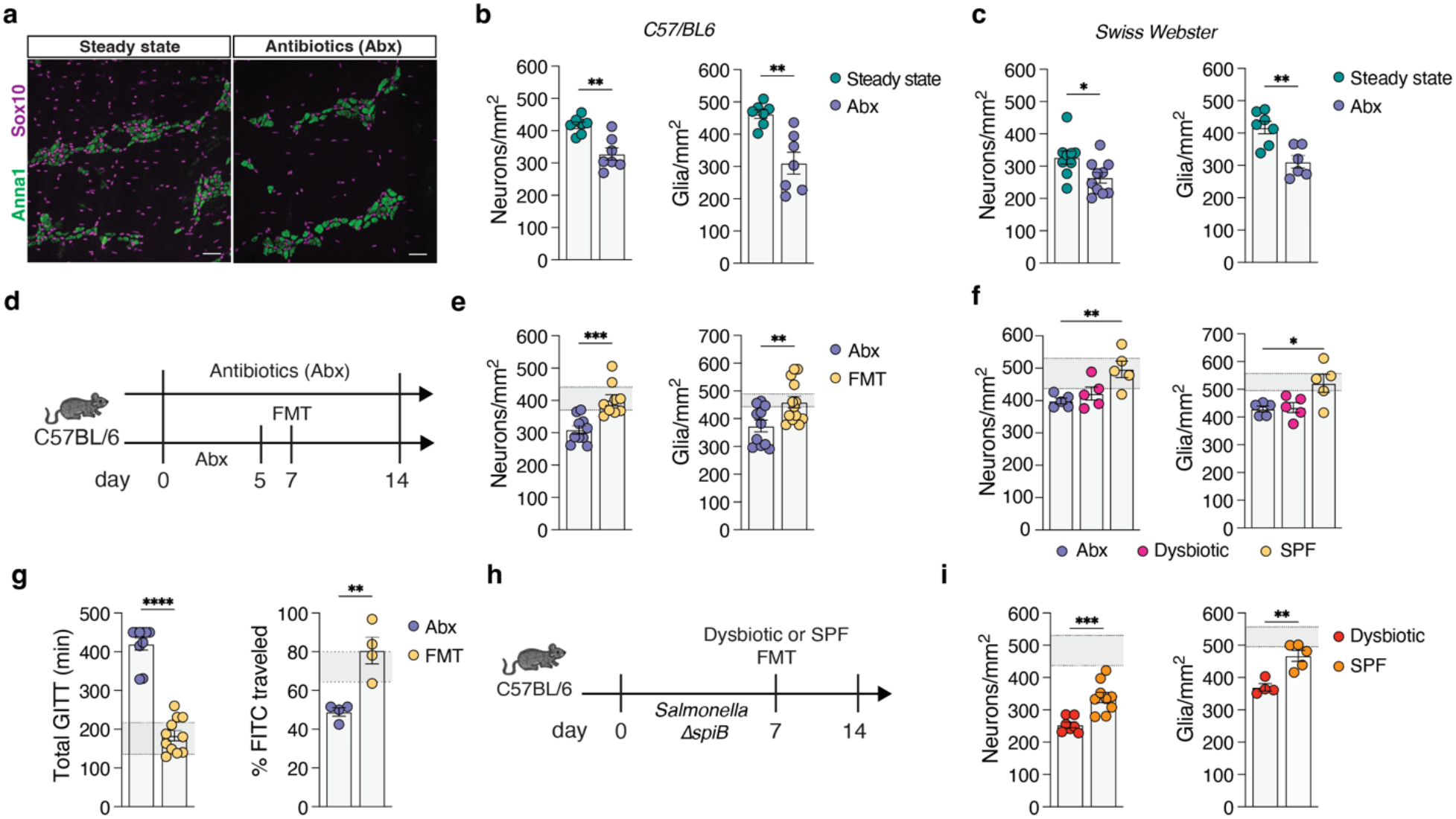
Microbiota-dependent maintenance of enteric neurons and glia. **a**, Representative confocal microscopy images of the mouse ileum myenteric plexus at steady state and five days post-Abx treatment, immunostained for Anna1 and Sox10. Scale bars, 50 μm. **b,c,** Quantification of enteric neurons (Anna1) and intraganglionic glia (Sox10) in C57BL/6 (**b**) and Swiss Webster (**c**) mice at steady state or following five days of Abx treatment. n = 7,7 (**b**); n = 9,10 (neurons) and 7,6 (glia) (**c**). **d,** Schematic representation of fecal microbiota transfer (FMT) following Abx treatment. **e,f,** Quantification of enteric neurons and intraganglionic glia 7 days after FMT. In-house C57BL/6 mice received continuous Abx treatment or FMT from SPF donors (**e**); Jackson C57BL/6J mice received FMT from dysbiotic or SPF donors following Abx treatment (**f**). The gray-shaded area in (e) indicates the range of neuronal and glial cell numbers at steady state in in-house C57BL/6 mice. n = 10,9 (neurons) and 11,13 (glia) (**e**); n = 5 (**f**). **g,** Effects of fecal microbiota transfer (FMT) following microbiota perturbation. Total gastrointestinal transit time (left) and small intestinal transit time (%FITC traveled, right) were measured in antibiotic-treated SPF mice seven days after FMT (**g**). **h,** Schematic representation of FMT following *Salmonella ΔspiB* infection. **i,** Quantification of enteric neurons and intraganglionic glia in *Salmonella ΔspiB*-infected C57BL/6J mice seven days after FMT from dysbiotic or SPF donors. The gray-shaded areas indicate the steady-state range for transit time (**g**) and neuronal/glial cell numbers in C57BL/6J mice (**i**; see **f**). (n = 10,11 (total GITT) and 4,4 (SITT) (**g**); n = 7,9 (neurons) and 4,5 (glia) (**i**). All data are presented as mean ±SEM. An unpaired two-tailed Student’s t-test was used for panels **b, c, e**, **f**, and **i.** One-way ANOVA, followed by Dunnett’s multiple comparisons test, was used for **g.** All data were obtained from the ileal myenteric plexus. Data are from at least two independent experiments, except for the %FITC traveled graph in panel **g** and glial quantification in panel **i.**

We observed that the total length of the small intestine changes with Abx treatment and returns to its steady-state length upon fecal transfer (**Extended Data Fig. 1b**). This raises the question of whether an increase in the total length of the small intestine contributes to a decrease in neuronal density. To assess this possibility, we quantified the number of neurons and glia per ganglion in all conditions. The average number of neurons or glia per ganglion decreased with Abx treatment and recovered upon SPF FMT, but not with dysbiotic FMT (**Extended Data Fig. 1c**), indicating that the change in neuronal density is not primarily driven by changes in the small intestinal length. To further rule out that the observed decrease in neuronal numbers upon Abx treatment was not due to structural changes, such as tissue stretching, we measured the average cell-to-cell distance between cell bodies within the ganglia. We did not observe any differences between the conditions (**Extended Data Fig. 1d**). Neuronal loss was independently confirmed in Baf53-Cre^INTACT^ mice, which express super-folder GFP fused to a nuclear membrane protein in the neuronal lineage^18^, ruling out possible artifacts in the quantification of neurons due to changes in the expression of the neuronal marker Anna1 (**Extended Data Fig. 1e**). We also did not observe a significant change in either cell number or small intestine length in GF mice treated with Abx, suggesting that the effects of Abx are not due to its potential cellular toxicity (**Extended Data Fig. 1f, g**). Finally, we tested whether reversing dysbiosis induced by *Salmonella* Δ*spiB* infection, which causes long-term loss of enteric neurons^2^, would also rescue neuronal and glial cell numbers (**Fig. 1h**). Dysbiotic FMT from mice during *Salmonella* infection did not rescue *Salmonella*-induced neuronal and glial loss; however, SPF FMT partially rescued both neurons and glial cells (**Fig. 1i**). These results indicate that enteric neuron and glial loss caused by microbial dysbiosis is reversible upon signals from healthy microbiota.

### Microbiota-derived succinate induces enteric neuron recovery in a tuft-cell-dependent manner

GF mice have fewer enteric neurons in the ileum^3^, providing a reductionist experimental platform for identifying microbial species and signals sufficient to induce ENS recovery. Therefore, we asked whether defined microbial communities are sufficient to recapitulate the increase in enteric neurons observed following colonization of GF mice with fecal microbiota from SPF mice^3,19^. We first compared enteric neuron numbers in GF mice and mice harboring the vertically transmitted, defined 12-member microbial consortium Oligo-MM12, which mimics key features of the mouse microbiota^20^ (**Fig. 2a**). Although vertically colonized Oligo-MM12 mice exhibited only a modest increase in enteric neuron number, GF mice vertically colonized with a supplemented consortium containing additional three facultative anaerobes^20^ (MM12+3) showed a robust increase (**Fig. 2a**).

**Fig. 2:**
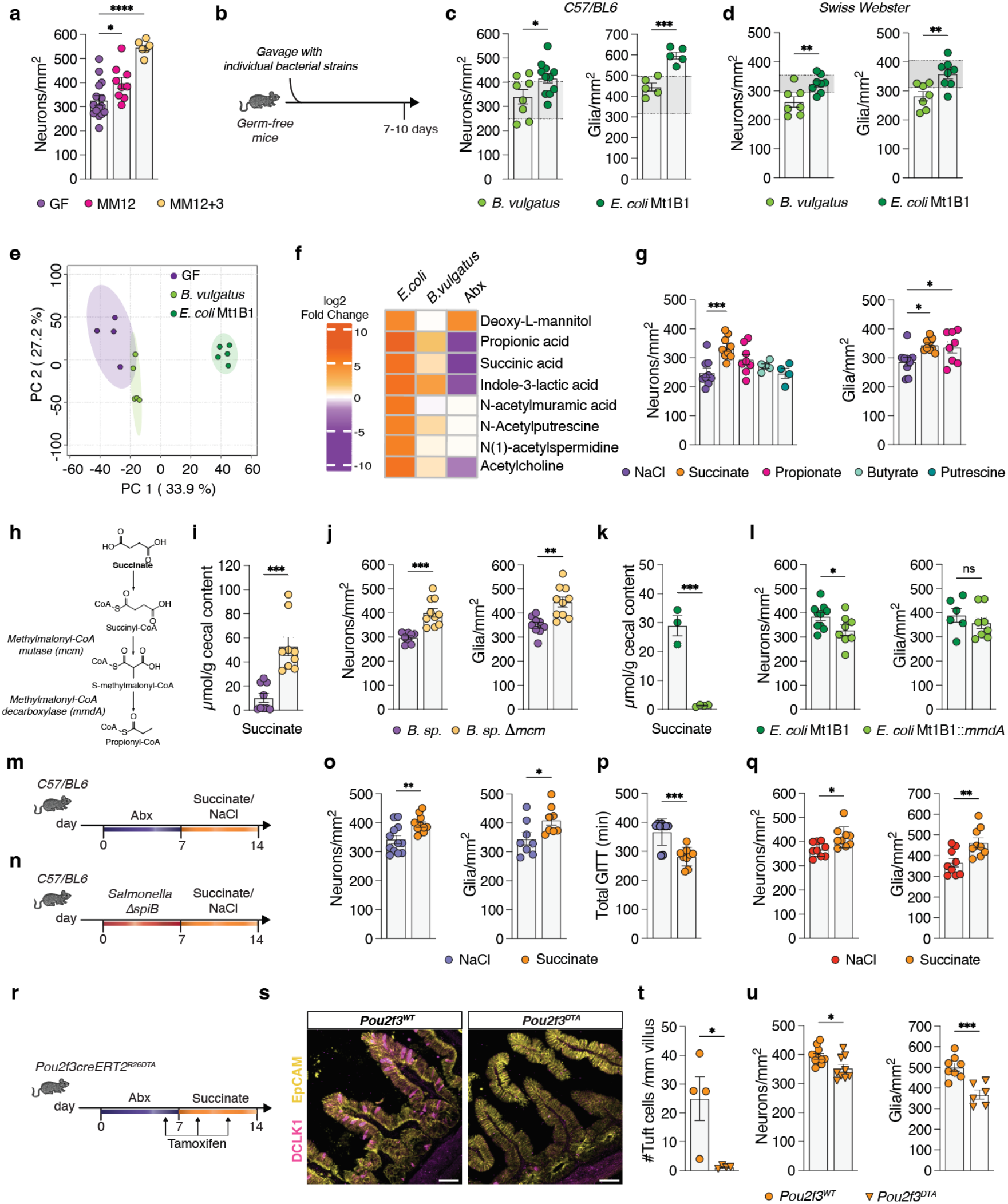
The gut bacteria-derived metabolite succinate is necessary and sufficient to induce enteric neuron recovery in a tuft cell-dependent manner. **a**, Quantification of neurons per mm² in the ileal muscularis of germ-free (GF) mice and GF mice colonized with defined microbial communities. *n =* 15, 9, 6 (bars from left to right). **b,** Schematic representation of GF mice monocolonization experiment. **c, d** Quantification of enteric neurons (left) and intraganglionic glia (right) in the ileum of GF C57BL/6 (**c**) and Swiss Webster (**d**) mice monocolonized with *B. vulgatus* or *E. coli* Mt1B1 for seven–ten days. Gray shading indicates the baseline neuronal and glial cell numbers in GF mice. *n =* 8, 11 (neurons) and *n* = 5, 5 (glia) for (c); *n =* 7,8 for (**d**). **e,f,** Untargeted cecal metabolomics following monocolonization. Principal component analysis of cecal metabolites from C57BL/6 GF mice and GF mice monocolonized with *B. vulgatus* or *E. coli* Mt1B1 for seven days (**e**), and selected metabolite heatmap including SPF mice treated with antibiotics for five days (**f**). n = 5 mice per group. **g,** Quantification of neurons (left) and intraganglionic glia (right) per mm^2^ area in Swiss Webster GF mice treated with metabolites in drinking water for seven days *n =* 9,9,8,5,4 bars left to right (neurons), *n=*9,9,8 (glia). **h,** Schematic representation of the methylmalonyl-CoA mutase pathway in *Bacteroides*. **i–l**, GF mice were monocolonized with the indicated bacterial strains for seven–ten days. Cecal succinate levels were quantified using HPLC-MS (**i**, **k**), and enteric neurons (left) and intraganglionic glia (right) were quantified per mm² (**j**, **l**). *n =* 5 (*i*); n = 10,10 (**j**); n = 3,4 (**k**); *n =* 10,8 (neurons), and 9,9 (glia) (*l*). **m, n,** Schematic representation of succinate rescue experiments in previously Abx (**m**) treated or *Salmonella ΔspiB* (**n**) infected mice. **o–q** Effects of succinate supplementation following microbiota perturbation. Quantification of enteric neurons (left) and intraganglionic glia (right) per mm² in Abx-treated mice receiving NaCl or succinate in drinking water for seven days (**o**) and in *Salmonella ΔspiB*-infected mice receiving NaCl or succinate for seven days (**q**). Intestinal transit time was measured in Abx-treated mice after seven days of NaCl or succinate supplementation (**p**). n = 12,11 (neurons) and 8,8 (glia) (**o**); n = 9,9 (**p**); n = 9,9 (**q**) **s,** Representative images of the intestinal epithelium in tuft cell-ablated and control mice immunostained with epithelial and tuft cell markers EpCAM (yellow) and DCLK1(magenta), respectively. **t,** Quantification of tuft cells (DCLK1+) per mm of crypt-villus axis. *n=* 4,4. Quantification of neurons (left) and intraganglionic glia (right) per mm^2^ area in tuft cell-ablated (Tx-treated) or control mice during succinate-mediated recovery for seven days. *n=*7,9 (neurons); *n=*8,6 (glia). All data are presented as mean ±s.e.m. An unpaired two-tailed Student’s t-test was used for panels **b, c, d, i, j, k, l,o, p, q, t,** and **u.** A one-way ANOVA, followed by Dunnett’s multiple comparisons test, was used for **g**. Data were obtained from at least two independent experiments.

We reasoned that one of these facultative anaerobes could mediate the observed increase in neuronal numbers. To test whether monocolonization with a single bacterial species, including one of the facultative anaerobes in MM12+3, was sufficient to increase the number of neurons and glia, we isolated several individual bacterial strains from the MM12+3 consortium and in-house SPF mice. Of those tested, the *Escherichia coli* Mt1B1 (*E. coli* Mt1B1) strain was the most potent in increasing both neuronal and glial numbers upon monocolonization in GF mice (**Fig. 2b, c,** and **Extended Data Fig. 2a, b**). Monocolonization with *E. coli* Mt1B1 also increased the number of neurons and glia in outbred Swiss Webster mice, suggesting that the observed neurogenic effect is robust and not confined to an inbred genetic background (**Fig. 2d**).

To identify the molecular signals mediating the observed increase in enteric neurons and glia, we focused on *E. coli* Mt1B1 and *Bacteroides vulgatus* (*B. vulgatus*) bacterial strains as reductionist models of “neurogenic” and “non-neurogenic” commensal bacteria, respectively. *Bacteroides* and *Escherichia* species differ in the immunogenicity of their lipopolysaccharide (LPS), with *E. coli* LPS being more immunostimulatory than *B. vulgatus*^21^. We tested whether the structural differences between the two commensals were sufficient to increase the number of enteric neurons and glia by treating GF mice with immunogenic LPS isolated from *E. coli*. We gavaged GF mice with *E. coli* LPS and observed no differences in the numbers of neurons and glia (**Extended Data Fig. 2c**). This prompted us to investigate the signals secreted by these two commensals. We performed untargeted liquid chromatography-mass spectrometry (LC-MS) analysis of polar metabolites in cecal contents isolated from mice monocolonized with *E. coli* Mt1B1 and *B. vulgatus* (**Supplementary Table 1**). The metabolomic profile of mice monocolonized with *B. vulgatus* was similar to that of GF mice, whereas mice monocolonized *with E. coli* Mt1B1 differed in the PC1 component in the principal component analysis (PCA), mirroring the ENS quantification phenotype (**Fig. 2e**). Among these candidates, short- and medium-chain fatty acids, such as propionic acid and succinate, which are the end products of bacterial fermentation, were enriched upon *E. coli* monocolonization (**Fig. 2f**).

Short-chain fatty acids (SCFAs) influence a wide range of host physiological processes, including immune cell differentiation, gut motility, neuronal activity, and the neurochemical code of enteric neurons^19,22–25^. We also observed an increase in polyamine-related metabolites, such as N-acetylputrescine and N-acetylspermidine, which we have previously shown to be neuroprotective against enteric infection-induced neuronal loss^2^. To test whether these metabolites could increase neuronal numbers, we treated GF mice with metabolites in drinking water *ad libitum*. Succinate treatment increased both neuronal and glial cell numbers in GF mice, whereas propionate had a modest gliogenic effect (**Fig. 2g**). We noted that succinate levels were also increased in the FMT state, as opposed to Abx and dysbiotic FMT (**Extended Data Fig. 2d**). Next, we orthogonally validated the effect of succinate by monocolonizing GF mice with the commensal protist *Tritrichomonas musculis* (*T. musculis*), which excretes high levels of succinate upon colonization in GF mice^26^. Consistent with succinate administration in drinking water, colonization with *T. musculis* also increased the number of enteric neurons and glia in GF mice (**Extended Data Fig. 2e**). Succinate is an intermediate metabolite generated during the fermentation of dietary fibers into SCFAs by gut commensals and provides critical energy substrates for host and microbial cells^27^. As a gain-of-function strategy to increase luminal succinate levels in “non-neurogenic” *Bacteroides* species, we generated a *Bacteroides* sp. 1_1_6 mutant targeting the methylmalonyl-CoA mutase gene (*Bacteroides sp.* 1_1_6 Δ*mcm*) using the allelic exchange strategy^28^. When CO_2_ levels are limited, *Bacteroides* species can regenerate CO_2_ via methylmalonyl-CoA mutase, which isomerizes succinate to methylmalonate; methylmalonate is then decarboxylated to generate CO_2_ and propionate^28,29^ (**Fig. 2h**). Disruption of the methylmalonyl-CoA mutase gene in *Bacteroides sp.* 1_1_6 *Δm*cm (*B. sp. Δmcm*) impaired succinate metabolism, resulted in elevated luminal succinate levels and a robust increase in neuronal and glial numbers following monocolonization of GF mice (**Fig. 2i, j**). As a complementary loss-of-function strategy to decrease bacterial succinate levels, we engineered the neurogenic *E. coli* Mt1B1 strain to heterologously express the *Bacteroides mmdA gene,* a component of the methylmalonyl-CoA mutase pathway, thereby enhancing succinate metabolism and reducing luminal succinate levels. GF mice monocolonized with *E. coli* Mt1B1::*mmdA* mutant showed reduced succinate levels in their lumen compared to mice colonized with wild-type *E. coli* Mt1B1 (**Fig. 2k**). In contrast to the parental strain, *E. coli* Mt1B1::*mmdA* failed to increase neuronal numbers; however, glial recovery was less affected (**Fig. 2l**). These findings indicate that, in our gnotobiotic system, bacterial succinate is necessary and sufficient for neuronal recovery or expansion in the adult murine ENS, but is not strictly required for glial recovery, suggesting the existence of additional microbiota-derived pathways that regulate gliogenesis.

To test whether increased succinate levels could also rescue neuronal and glial loss induced by microbial dysbiosis upon Abx treatment or *Salmonella* Δ*spiB* infection, we administered succinate to SPF mice previously treated with Abx or infected with *Salmonella* Δ*spiB* (**Fig. 2m, n**). Both neuron and glial cell numbers were restored to their pre-infection levels, further validating the neurogenic and gliogenic properties of succinate (**Fig. 2o, q**). Succinate also rescued the increased transit time defect in Abx-treated mice and did not alter intestinal barrier permeability to either 4 kDa or 70 kDa FITC, suggesting that ENS recovery occurred in the absence of overt epithelial barrier disruption (**Fig. 2p** and **Extended Data Fig. 2f**). Succinate is sensed by tuft cells in the epithelium via its G protein-coupled receptor, GPR91 (SUCNR1), resulting in tuft cell hyperplasia and the induction of a type 2 immune response^30–32^. We investigated whether tuft cells mediate the effects of succinate on the recovery of the enteric nervous system following microbial dysbiosis. To directly address the requirement of tuft cells during succinate-mediated recovery after Abx treatment, we ablated tuft cells using tamoxifen-inducible tuft-cell-specific driver Pou2f3^creERT2^ crossed to R26^DTA^ mice (**Fig. 2r**). We confirmed the ablation efficiency using tuft cell-specific Dclk1 immunostaining in the epithelium (**Fig. 2s, t**). Tuft-cell-ablated mice failed to recover enteric neurons and glia following succinate treatment (**Fig. 2u**), indicating that exogenous succinate-mediated ENS recovery after Abx-induced microbial dysbiosis requires tuft cells. Together, these findings identify a succinate–tuft cell signaling axis that promotes recovery of the enteric nervous system following dysbiosis.

### Identification of ENS recovery and stress/injury-associated states upon microbial manipulations

The ENS develops from vagal and sacral neural crest-derived progenitors that migrate along the developing gut, giving rise to neurons and glia with a highly coordinated pattern of self-renewal and differentiation^9^. Although postnatal neurogenesis in the ENS ceases around weaning age, our observations suggest that the adult ENS retains mechanisms that promote neuronal recovery following microbiota perturbations. One possibility raised in previous studies is glia-to-neuron differentiation in chemical injury or colitis models^14,33^. Another is the continuous turnover of adult ENS from Nestin-expressing progenitor cells^34,35^. Therefore, we sought to identify the cellular populations associated with neuronal recovery following FMT. We established a single-nucleus RNA sequencing (snRNA-seq) workklow for the mouse ileal muscularis to enable the unbiased characterization of ENS cell populations across different microbiota conditions. To capture early cellular states associated with ENS recovery, we isolated nuclei from the ileal muscularis of mice at steady state, Abx, and three days post-FMT conditions and prokiled approximately 20,000 nuclei per condition using the 10x Genomics platform. Mesenchymal populations were identikied using canonical markers (*Kit*, *Myh11*) and excluded (**Extended Data Fig. 3a-c**). Nuclei with high mitochondrial content were also excluded (**Extended Data Fig. 3b, c**).

To test whether the proportion of ENS-associated clusters changed across treatments, we integrated the subsetted conditions, reclustered, and visualized them using UMAP (**Fig. 3a**). Cells with neuronal expression signatures were separated into six clusters, and those with glial signatures were divided into three clusters based on the expression of marker genes for specikic cell types (**Fig. 3c, d**). We observed signikicant changes in cluster abundance across conditions using Fisher’s exact test (**Extended Data Fig. 3d**). Cluster 5 (c5) was enriched following Abx-mediated perturbation of the microbiota and preferentially expressed genes associated with cellular stress and injury (*Gsdme*, *Mcu*, *and Eps8*) (**Fig. 3a-c**). We also identikied a small recovery-associated cluster with neuronal features and enrichment of cell cycle genes, which we annotated as a precursor-like state (cluster 11, c11). This population was enriched in Abx and FMT conditions and expressed cycling-associated genes, including *Top2a* and *Mki67* (**Fig. 3b, c,** and **Extended Data Fig. 4c, d**). Two clusters (c0 and c12) expressed Kit and mesenchymal markers and exhibited features consistent with those of the interstitial cells of Cajal (Fig. 3c).

**Fig. 3:**
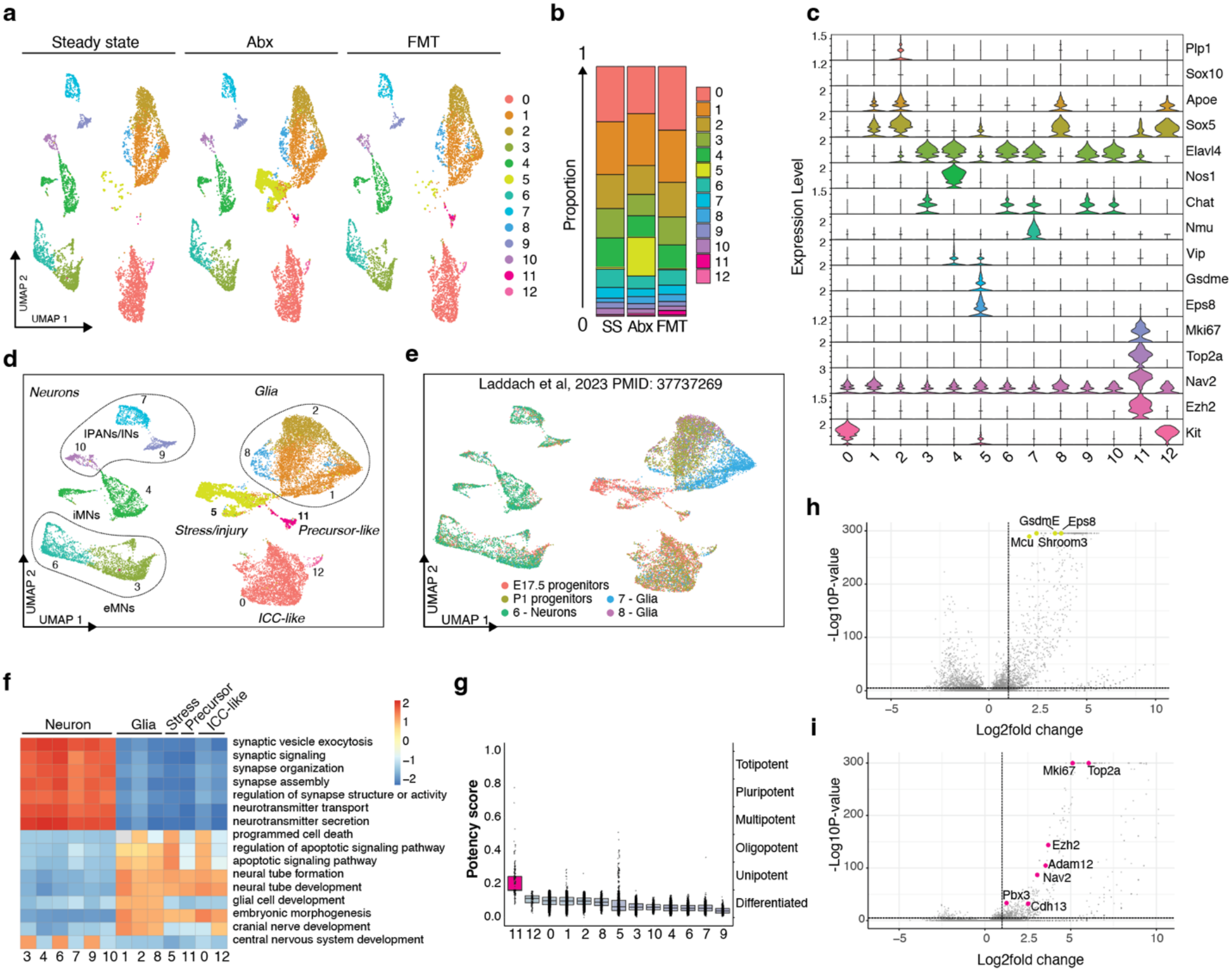
Single-nucleus RNA-seq (snRNA-seq) proJiling of myenteric plexus cell populations reveals ENS recovery and injury/stress-associated states. a, Split UMAP plot of subsetted datasets of muscularis layer nuclei from steady-state, Abx, and FMT mice after removal of mesenchyme (n=4 mice per condition, pooled). b, Bar plot displaying the frequency of each cluster under the individual conditions. c, Violin plot displaying the expression of selected marker genes in the clusters. d, UMAP plot of subsetted cells merged across all conditions with cluster annotation. iMN, inhibitory motor neurons; eMN, excitatory motor neurons; IPAN, intrinsic primary afferent neurons; IN, interneurons; ICC, interstitial cells of Cajal. e, UMAP plot with labels transferred from Laddach *et al*., 2023 scRNA-seq. f, Gene set enrichment analysis (GSEA) of the annotated clusters. g, *CytoTrace2* cell potency scores for the individual clusters. h, Volcano plot of genes differentially expressed in the stress/injury cluster (c5) compared to the other clusters. *log2foldchange cut-off =* 1, displayed as a dashed line. i, Volcano plot of genes differentially expressed in the precursor-like cluster (c11) compared to the rest of the clusters. *log2foldchange cut-off =* 1, displayed as a dashed line.

To further validate our annotations, we performed label transfer using published ENS single-cell atlases^36–38^. This analysis recovered the published enteric neuronal subtype identities within neuronal clusters (**Extended Data Fig 4a**), and the precursor-like cluster was mapped predominantly to embryonic (E17.5) and early postnatal (P1) ENS progenitor populations (**Fig. 3e**). Gene set enrichment analysis revealed that synaptogenesis and neurotransmitter release-related pathways were enriched in the neuronal clusters (**Fig. 3f**). In contrast, pathways associated with programmed cell death were enriched in the stress/injury cluster specikic to the Abx state (c5), and pathways related to neuronal development were enriched in the precursor-like cluster (c11) as well as glial clusters (**Fig. 3f**). To characterize the differentiation state of these populations, we applied *CytoTrace*, which infers relative developmental potential from transcriptional diversity^39^. The precursor-like cluster (c11) exhibited the highest developmental potency score among all populations classikied as “unipotent”; meanwhile, the remaining clusters were predominantly classikied as “differentiated” (**Fig. 3g**). These kindings suggest that c11 represents a relatively immature state associated with neuronal recovery.

Next, we searched for differentially enriched genes within the stress/injury and precursor-like clusters (**Supplementary Tables 2 and 3**). Among the differentially expressed genes, the stress/injury cluster included *Gsdme* and *Mcu*, both of which are implicated in mitochondrial damage that leads to neuronal death^34,35^ (**Fig. 3h**). To determine whether Abx-induced ENS loss depends on Gsdme, we treated Gsdme knockout mice with Abx and quantikied the number of enteric neurons and glia. In contrast to Abx-treated wild-type mice, Gsdme-dekicient mice were protected from both neuronal and glial loss following Abx treatment, indicating that Gsdme contributes to ENS loss in Abx-induced microbiota perturbation (**Extended Data Fig. 4c**).

The precursor-like cluster was enriched in cell cycle-related genes such as *Top2a*, the chromatin regulator *Ezh2*, and the retinoic acid-responsive gene *Nav2,* which is involved in neuronal migration and neurogenesis in the central nervous system (CNS)^40^ (**Fig. 3i and Extended Data Fig. 4d, e**). Since *Nav2* was selectively enriched in the precursor-like cluster and associated with microbiota-dependent recovery, we selected it as a marker to further investigate recovery-associated cellular states in the adult ENS.

### Plp1+ glia acquire recovery-associated cellular states following Abx treatment

Next, we sought to determine whether Nav2-associated cellular states emerge during ENS recovery following FMT. Pseudotime trajectory inference positioned the precursor-like cluster at the root of the inferred differentiation trajectory, suggesting that it may represent an early recovery-associated state (**Fig. 4a**). Consistent with this interpretation, *Nav2* expression was detected in proliferative ENS precursor populations during postnatal development on day seven, supporting its association with immature neuronal states, coinciding with the expression of the glial marker *Plp1*^41^ (**Fig. 4b, c**). To investigate the relationship between Nav2-associated recovery states and the Plp1 lineage, which has been postulated to differentiate into neurons following colitis and, to a lesser extent, following chemical injury^14,33^, we combined chromatin profiling of Plp1+ glia with genetic fate-mapping approaches.

**Figure 4:**
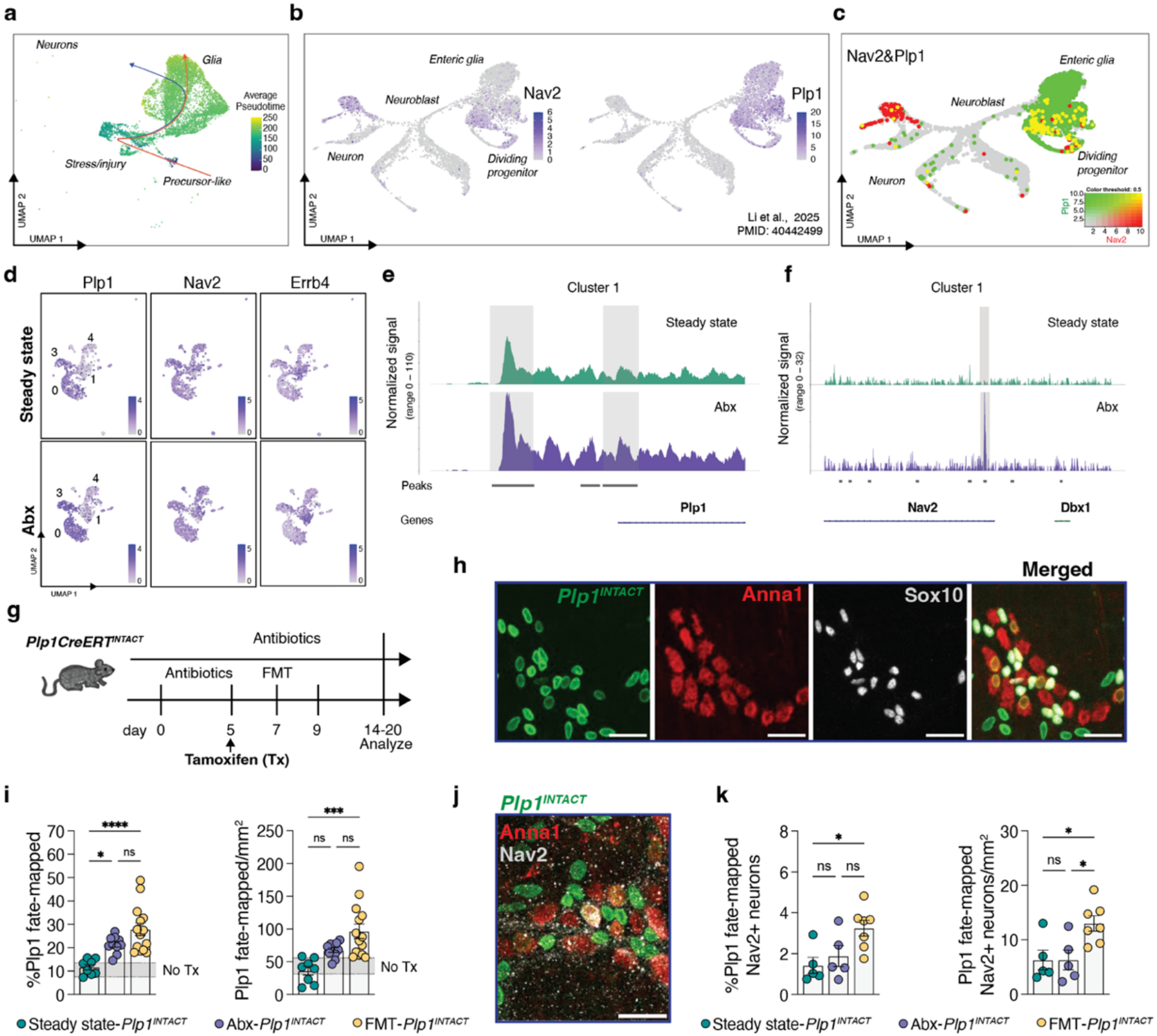
Plp1+ glia acquire a Nav2-associated recovery state during microbiota-dependent ENS recovery. **a,** Pseudotime trajectory inference using Slingshot on precursor-like, stress/injury, and glial clusters of the snRNA-seq dataset from Fig. 3. **b,** Feature plot displaying Nav2 and Plp1 in the postnatal d7 ENS scRNA-seq dataset by Li *et al*., 2025. Cluster annotations were obtained from the original publications. **c,** Feature plot displaying cells that co-express Nav2 and Plp1 in the postnatal d7 ENS scRNA-seq dataset in Li *et al*., 2025. Cluster annotations were obtained from the original publication. *Color-threshold = 0.5*. **d,** Gene activity scores for *Plp1*, *Nav2*, and *Errb4* in scATAC- seq profiles of Plp1+ cells from steady-state and Abx-treated mice (n = 5 mice per condition, pooled). **e,f,** Genome browser tracks showing normalized chromatin accessibility at the *Plp1* (**e**) and *Nav2* (**f**) loci. The gray-shaded area indicates differentially accessible peaks between the steady-state and Abx-treated conditions. **g,** Schematic representation of the experimental design for the inducible Plp1 glial fate-mapping experiment. **h,** Representative confocal microscopy images of the ileal myenteric plexus of *Plp1^INTACT^* (green) inducible reporter mice immunostained with AnnaA1 and Sox10. Scale bars, 50 μm. **i,** Quantification of the percentage of *Plp1^INTACT^*+Anna1+ cells among total Anna1+ neurons (left) and the density of *Plp1^INTACT^*+Anna1+ cells per mm^2^ of ileal myenteric plexus (right) seven–ten days following Abx or FMT. The gray-shaded area indicates the range of neuronal densities in mice in the steady state. *n =* 8,11,14 (bars from left to right). **j,** Representative confocal image showing a *Plp1^INTACT^* + labeled cell immunostained for Anna1 and Nav2 proteins. Scale bar, 50 μm. **k,** Quantification of the percentage of *Plp1^INTACT^*+Anna1+Nav2+ cells among total Anna1+ neurons (left) and the density of *Plp1^INTACT^*+Anna1+Nav2+ neurons per mm^2^of the ileal myenteric plexus (right) seven–ten days following Abx or FMT. *n =* 5,5,7 (bars from left to right). All data are mean ±s.e.m. A one-way ANOVA, followed by Tukey’s multiple comparisons test, was used for **i** and **k**.

We generated Plp1^creERT^; INTACT mice and analyzed chromatin accessibility in Plp1+ glia following Abx treatment using the 10x Genomics Epi ATAC single-cell ATAC-seq (scATAC-seq) platform (**Extended Data Fig. 5a-d**). We recovered 11 distinct clusters and performed label transfer using previously published p7 and p21 ENS scRNA-seq datasets to further define their molecular identities (**Extended Data Fig. 5d-f**). Clusters 1 and 4 acquired neuronal annotations, while cluster 1 also aligned with p7-dividing progenitor, enteric glia, and neuroblast populations, suggesting the acquisition of immature developmental features (**Extended Data Fig. 5e, f**). Moreover, Abx-treated glia exhibited increased accessibility at the *Nav2* locus, especially in cluster 1, which also showed elevated activity of the ENS progenitor-associated gene *Errb4*, consistent with activation of a neuronal recovery-associated program with developmental features (**Fig 4d-f**). Among the significantly enriched peaks in this cluster were the *Plp1* upstream regulatory region and the 3′ end of the *Nav2* locus, labeled across the gray shaded line (**Fig. 4e, f, and Extended Data Fig. 5g and h; Supplementary Table 4**). Together, these findings suggest that the *Nav2* locus becomes poised for activation in a subset of Plp1+ cells following microbiota perturbation.

To fate-map Plp1+ cells during FMT-mediated recovery, we administered tamoxifen (Tx) on the last day of Abx treatment and traced whether the labeled Plp1+ cells acquired a neuronal identity using immunohistochemistry coupled with confocal imaging (**Fig. 4g, h**). We observed approximately 10% Plp1+ neurons in the absence of Tx, indicating the leakiness of the inducible driver (**Fig. 4h**). The 10% misexpression rate in neurons was similar to that of the Tx-only control, suggesting little or no glia-to-neuron differentiation or turnover at steady state (**Fig. 4h**), which is consistent with previous studies suggesting limited neurogenesis in the adult ENS under steady-state conditions^13,15,42^.

In contrast, the percentage of fate-mapped neurons increased significantly to approximately 25% following Abx treatment and FMT-mediated recovery (**Fig. 4h**). To distinguish the contribution of pre-existing Plp1+ cells from the possibility that Plp1 expression was induced during Abx treatment, we performed an independent fate-mapping experiment in which Plp1+ cells were labeled with Tx before Abx exposure. The number of fate-mapped neurons increased following Abx treatment under this pre-Abx labeling scheme, relative to post-Abx labeling, supporting the contribution of pre-existing Plp1+ cells rather than the induction of Plp1 expression during Abx treatment (**Extended Data Fig. 6a**). Moreover, both the percentage and absolute number of Nav2+ Plp1+ fate-mapped neurons increased during FMT, consistent with the activation of a Nav2-associated recovery program within the Plp1+ lineage (**Fig. 4j, k**). While we observed increased fate mapping upon Abx treatment alone, the net neuronal numbers only recovered following FMT (**Fig. 1e, g**). Together with the changes in chromatin accessibility in Plp1+ cells upon Abx treatment, these findings support a model in which microbiota depletion primes a Plp1-associated recovery program, whereas microbiota restoration promotes the induction or maintenance of recovery-associated neurons. To further investigate the cellular dynamics underlying this response, we assessed the proliferation of Plp1+ cells using the thymidine analog 5-ethynyl-2’-deoxyuridine (EdU). The fraction of EdU+ Plp1+ cells was low (∼0.4%) and did not differ between the Abx and FMT conditions, as determined by flow cytometry (**Extended Data Fig. 6b**). These findings suggest that the activation of the Plp1-associated recovery program occurs largely independently of substantial glial proliferation.

Next, we investigated whether this recovery response was specific to the Plp1 lineage or was shared across other enteric glial populations. Growing evidence suggests heterogeneity in enteric glial cells at the molecular, chromatin, and functional levels^43–45^. To determine whether glial populations other than Plp1+ cells contribute to neuronal recovery, we performed complementary lineage-tracing experiments using Tx-inducible Sox10^creERT2^ and GFAP^creERT2^ drivers (**Extended Data Fig. 6c**). No significant changes in the percentage of Sox10+ or GFAP+ fate-mapped neurons were observed between the Abx, FMT, and steady-state groups (**Extended Data Fig. 6d, e**). Similar results were obtained using the independent SER93-Sox10^creERT2^ line, which shows minimal leaky expression^15^ (**Extended Data Fig. 6f**). These findings suggest that the recovery-associated response is not a general feature of all enteric glial populations but is preferentially associated with the Plp1 lineage. Taken together, our findings suggest that a subset of Plp1+ cells participates in a Nav2-associated recovery program in response to microbiota perturbations and contributes to neuronal recovery.

### Succinate activates developmental neuronal state transitions in the adult ENS

Having linked Plp1+ glia to ENS recovery in complex microbiota, we next asked whether a single, well-defined signal is sufficient to activate the neuronal program associated with this response. Therefore, we turned to a reductionist gnotobiotic model using monocolonization of GF mice with the succinate-accumulating mutant *B. sp. Δmcm* to determine whether a single, defined microbial signal can recapitulate the developmental neuronal state transitions in the adult ENS. Recent single-cell expression profiling studies have shown that ENS neuronal subtype specification follows stereotypical developmental trajectories, in which early branch populations progressively give rise to distinct neuronal subtype identities^37,41^. To determine whether succinate-induced neuronal recovery recapitulates aspects of this developmental framework, we rederived *Plp1CreERT^INTACT^* mice under GF conditions and monocolonized them with succinate-accumulating mutant *B. sp.* Δ*mcm.* We isolated nuclei from the ileal muscularis layer on day 0 (GF, before colonization), day five, and day 14 following monocolonization, representing the early and late stages of differentiation (**Fig. 5a and Extended Data Fig 7a, b**). Despite enrichment for GFP+ P*lp1^INTACT^* glial nuclei and Anna1+ neuronal nuclei, we recovered a small number of muscle and ICC populations, which were removed before downstream analyses (**Extended Data Fig. 7c-e**). We annotated neuronal populations by label transfer using the ENS subtype atlas generated by Morarach *et al.* and recovered most neuronal classes, except for ENC11 (**Fig. 5b**)^37^. Although GF mice exhibit reduced neuronal numbers (**Fig. 2b**), all major neuronal subtype classes were represented across conditions, indicating preservation of the overall molecular architecture of the ENS (**Extended Data Fig. 7f**).

**Fig 5.**
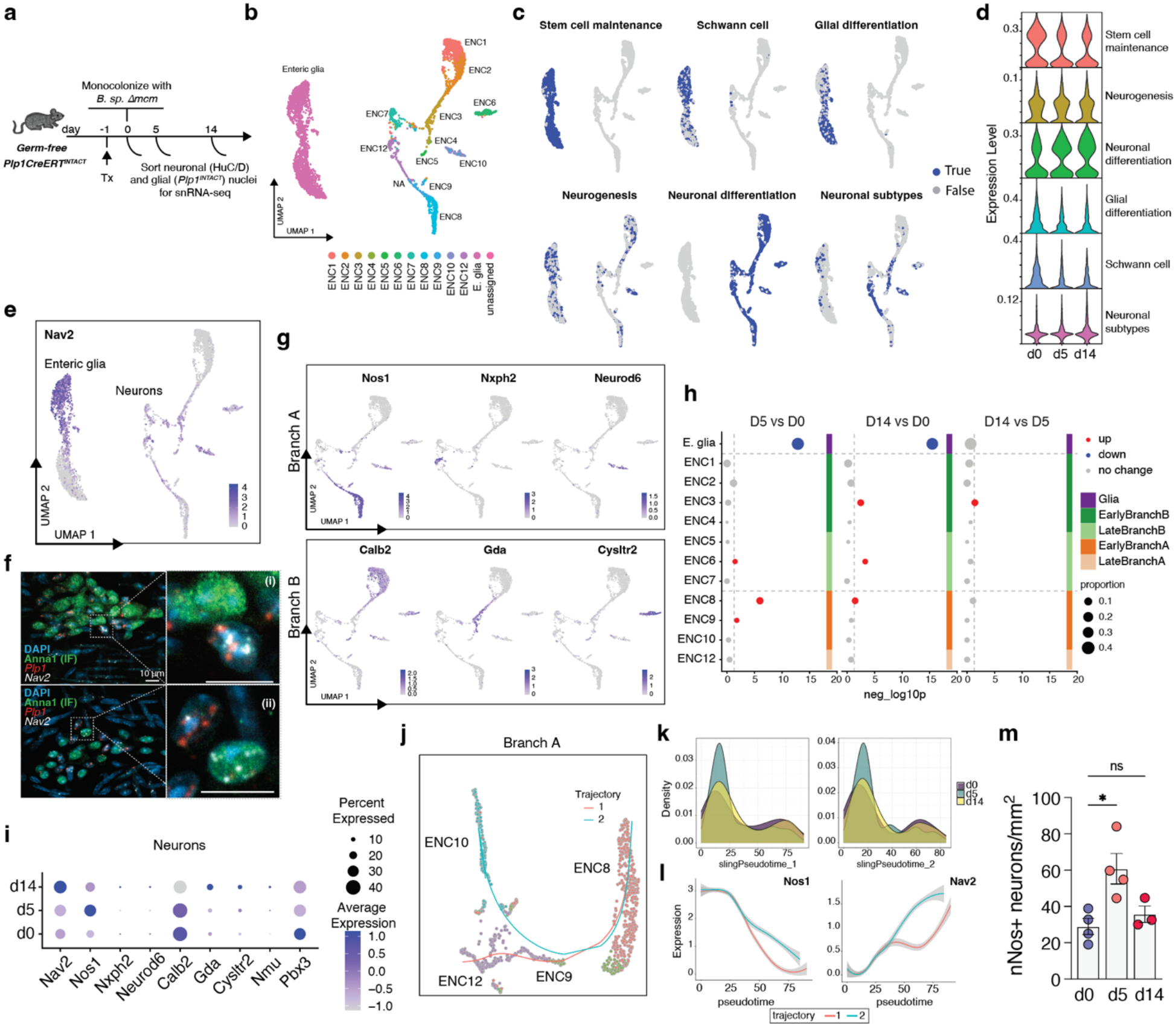
Succinate induces developmental-like neuronal state transitions and subtype diversification in the adult ENS. **a**, Experimental design for time-series snRNA-seq of the ENS following monocolonization of GF mice with *B. sp. Δmcm*. **b,** Merged UMAP plot annotated by label transfers using the reference dataset Morarach *et al*. (2020), showing the individual ENS subtypes. *N =* 4 mice per condition, pooled. **c,** UMAPs displaying gene set scores for neuronal subtype-specific gene signatures defined by Morarach *et al.* (2020). **d,** Violin plot showing the distribution of gene set scores in **c** across conditions. **e,** Feature plot showing the expression of *Nav2* in the merged snRNA-seq dataset. **f,** Representative RNAscope images showing *Plp1* and *Nav2* transcripts in **(i)** *Plp1*+*Nav2*+ glial cells. **(ii)** *Nav2*+Anna1+ neurons. Scale bars, 10 µm. **g,** Feature plots showing the expression of representative ENS subtype marker genes across Branch A and B neuronal populations, consistent with the developmental trajectories described by Morarach *et al.* (2020). **h,** Changes in the relative abundance of neuronal subtypes across experimental conditions within Branches A and B, assessed using Fisher’s exact test. **i,** Dot plot showing the expression of representative neuronal branch marker genes under different experimental conditions. **j,** Slingshot trajectory inference of Branch A neuronal populations. **k,** Density plots showing the distribution of cells along the Branch A-inferred trajectories shown in **j**. **l,** Expression of *Nos1* and *Nav2* along the trajectories shown in **j. m,** Quantification of Nos1+ neurons per mm² of the ileal muscularis in GF mice (d0) and in mice five- and 14-days following *B. sp. Δmcm* monocolonization. *n =* 4,4,3. One-way ANOVA followed by Dunnett’s multiple comparisons test.

Gene-set signature scoring revealed enrichment of stem-cell maintenance, Schwann-cell, and glial differentiation programs in the GF state, whereas neuronal differentiation and neuronal subtype specification programs increased following succinate exposure and were most prominent on day 14 (**Fig. 5c, d**; differential gene expression across conditions is provided in **Supplementary Table 5**). Consistent with these findings, *CytoTrace* analysis showed that the GF sample contained the highest proportion of oligopotent cells, whereas no oligopotent cells were detected in the succinate day 14 population, consistent with a shift toward a more differentiated state (**Extended Data Fig. 8a, b**). Among all populations, glial clusters displayed the highest developmental potential scores (**Extended Data Fig. 8c**). Consistent with earlier analyses, *Nav2* expression was detected in Schwann-like glial populations and subsets of differentiated neurons (**Fig 5c, e**, and **Extended Data Fig. 8d, f**). Although Nav2 protein expression was readily detected in neurons by immunohistochemistry, we did not observe consistent detection in glial cells. Therefore, we used fluorescent *in situ* hybridization (RNAscope) to independently validate *Nav2* transcript expression in both glial and neuronal populations (**Fig. 5f**).

Next, we investigated whether succinate-induced recovery recapitulates aspects of developmental ENS subtype diversification. Our dataset captured both major neuronal branches described by Morarach *et al.*, starting with inhibitory motor neuron ENC8 (Branch A) and excitatory motor neuron ENC1 (Branch B) lineages, allowing us to examine subtype dynamics over time (**Fig. 5g, Extended Data Fig. 8e**). Fisher’s exact tests revealed changes in the relative abundance of neuronal subtype populations across time points (**Fig. 5h**). Nos1+ ENC8 neurons, which occupy an early position within Branch A, were enriched on day five and declined on day 14. In contrast, neuronal populations associated with later developmental states, including ENC6 neurons within Branch B, expanded by day 14 (**Fig. 5h, i**). To visualize these temporal changes, we performed pseudotime analysis separately for Branches A and B. Cells from day five occupied earlier pseudotime positions, whereas day 14 cells were enriched toward terminal branch states (**Fig. 5j-l** and **Extended Data Fig. 8g-i**). Notably, *Nav2* expression increased toward the distal ends of both inferred trajectories, consistent with its association with recovery-associated neuronal states (**Fig. 5k, l,** and **Extended Data Fig. 8h, i**). Consistent with these analyses, Nos1+ neurons transiently increased on day five, as determined by immunohistochemistry (**Fig. 5m**). In addition to the changes in subtype composition, succinate exposure increased the expression of activity-dependent immediate early genes (IEGs), including *Fos*, *Fosb, Jun, Junb, Egr1, Nr4a1, Arc,* and *Npas4*, consistent with the activation of a neuronal transcriptional program associated with synaptic plasticity, potentially reflecting activity-dependent circuit remodeling (**Extended Data Fig. 8j**). Collectively, these findings suggest that succinate-induced neuronal recovery partially recapitulates the developmental diversification model of the ENS, characterized by the transient expansion of early ENC8-like neuronal states, followed by the emergence of late branches associated with distinct neuronal subtype identities. Thus, a single microbiota-derived signal is sufficient to promote developmental-like neuronal state transitions and molecular programs associated with activity-dependent circuit remodeling in the adult ENS.

### Succinate increases Nav2+ neurons through a Plp1-independent recovery pathway

Our lineage tracing indicated that a subset of Plp1+ cells contributes to neuronal recovery through a Nav2-associated recovery program initiated by microbiota perturbation. Therefore, we investigated whether succinate-mediated neuronal recovery is also driven by an increased contribution of Plp1+ cells. We first tested whether succinate contributes to Plp1+ neuron fate mapping using our Plp1-based fate-mapping approach (**Extended Data Fig. 9a**). However, we did not observe an increased number of Plp1-labeled neurons during succinate-mediated neuronal recovery in previously Abx-treated mice (**Extended Data Fig. 9a, b**), despite the recovery of neuronal numbers (**Fig. 2o**). Because Abx recovery is a dynamic process, we used a gnotobiotic setting in which *B. sp.* Δ*mcm* increases neuronal numbers without Abx-induced loss. To this end, we used the same monocolonization paradigm and fate-mapped Plp1+ cells following monocolonization with either WT *B.* sp. or the succinate-accumulating mutant *B. sp.* Δ*mcm*. Plp1+ cells were labeled with sterile Tx one day before colonization (**Fig. 6a**). Despite the increase in neuronal numbers induced by the succinate-accumulating mutant, we did not observe an increase in the number of Plp1 fate-mapped neurons (**Fig. 6b**). Together with the absence of injury-associated signals in the gnotobiotic setting, these findings suggest that Plp1-associated recovery is linked to microbiota perturbation, whereas succinate-induced neuronal accumulation proceeds via a distinct pathway. Finally, because succinate was sufficient to partially recapitulate developmental neuronal state transitions, we investigated whether it also expanded the Nav2-associated recovery state identified during FMT-mediated recovery (**Fig. 3**). Monocolonization with the succinate-accumulating mutant *B. sp. Δmcm* significantly increased both the percentage and total number of Nav2+ neurons compared to WT-colonized controls (**Fig. 6c, d**). Consistent with this observation, Nav2+ neurons were similarly enriched during succinate-mediated recovery following Abx treatment (**Fig. 6e**). Collectively, these findings suggest that microbiota-derived succinate promotes neuronal accumulation by expanding Nav2-associated neuronal states via a pathway distinct from the Plp1-associated recovery program.

**Fig. 6:**
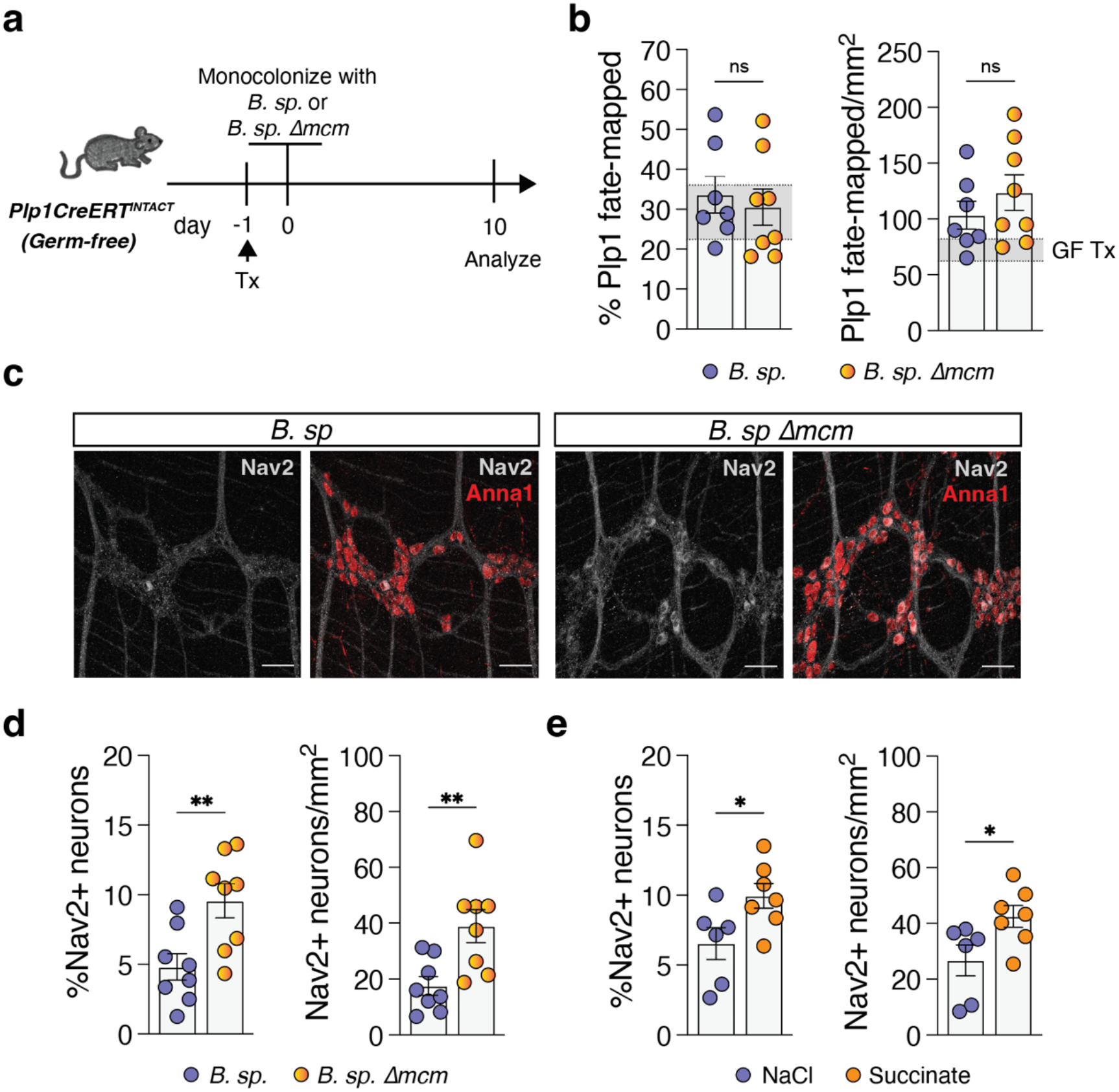
Succinate promotes Nav2-associated neuronal recovery independent of increased Plp1+ lineage contribution. **a,** Schematic representation of the experimental design of Plp1 glia fate-mapping experiment in GF mice monocolonized with *B. sp.* or *B. sp.* Δ*mcm*. **b,** Quantification of the percentage of *Plp1^INTACT^*+ Anna1+ fate-mapped cells among total Anna1+ neurons (left) and the density of *Plp1^INTACT^*+ Anna1+Nav2+ neurons per mm^2^ of ileal myenteric plexus (right) 10 days after monocolonizations. *n =* 7, 8 (bars left to right). The gray shaded region represents the baseline quantification in GF Tx-treated animals. **c,** Representative confocal microscopy images of Anna1 and Nav2 immunostaining after monocolonization. Scale bars, 50 µm. **d,** Quantification of the percentage of Anna1+ neurons among total neurons (left) and the density of Anna1+ neurons per mm2 of the ileal myenteric plexus (right) 10 days after monocolonization. *n =* 8, 8 (bars left to right) **e,** Quantification of the percentage of Anna1+ neurons among total neurons (left) and the density of Anna1+ neurons per mm2 of the ileal myenteric plexus (right) seven days after succinate or NaCl treatment following Abx in SPF mice *n =* 6, 7 (bars left to right). All data are mean ±s.e.m. A one-way ANOVA, followed by Tukey’s multiple comparison test, was used for **i and k**. An unpaired two-tailed Student’s t-test was used for **panels b, d, and e**.

## Discussion

This study identified gut bacterial species with neurogenic and non-neurogenic roles, and the bacterial metabolite succinate as a key downstream signal to regenerate enteric neurons and glia in adult mice. Neuronal and glial recovery exhibited distinct succinate dependencies. While succinate was necessary and sufficient for neuronal recovery or expansion, its role in glial recovery appeared more complex, as the loss of bacterial succinate production in a gnotobiotic setting did not result in a comparable defect in glial recovery. These findings suggest that enteric glial recovery can be mediated by multiple microbiota-derived signals, with succinate as one component of a broader recovery network not captured by reductionist gnotobiotic models.

Fate-mapping of enteric glial populations suggested that a Plp1-associated program contributes to neuronal recovery initiated during Abx treatment. Therefore, it is plausible that enteric glia are involved in tissue surveillance that detects damage and enables the rapid onset of regeneration upon injury, with replacement components at hand^46^. An intriguing aspect of our findings is that succinate appears to exert distinct effects depending on the physiological context. During recovery following Abx-induced dysbiosis, succinate was sufficient to restore neuronal numbers but did not increase Plp1 fate-mapping, suggesting that injury-associated signals initiate glial plasticity, whereas succinate may promote the survival and maturation of newly generated neurons. In contrast, under gnotobiotic conditions, succinate-accumulating *B. sp. Δmcm* mutant bacteria induced developmental neuronal state transitions and increased Nav2+ neurons without increasing Plp1-lineage contribution. Thus, succinate engages distinct recovery mechanisms depending on the tissue context. Consistent with this context dependence, pathogen-derived succinate exacerbates neuronal loss in an experimental colitis model^47^, whereas commensal-derived succinate promotes neuronal recovery in our model. These findings highlight that the effects of microbiota-derived metabolites are context-dependent and may be influenced by the underlying inflammatory state, tissue environment, and nature of the injury.

Growing evidence suggests that cellular plasticity occurs in enteric glia during adulthood. Recent studies have highlighted the neurogenic potential of enteric glial cells by retaining an open chromatin state at critical early ENS progenitor loci^38,43^. Our glial chromatin profiling and Plp1+ lineage-tracing results align with these recent observations and provide functional validation of the plasticity of enteric glial populations that respond to environmental stimuli. The limited EdU labeling in Plp1+ glia may reflect transdifferentiation without proliferation rather than dedifferentiation into a progenitor state^47^. A non-mutually exclusive possibility for the lack of EdU incorporation is a shortened S phase^48^, or cell division without new DNA synthesis involved^49^.

Although succinate can signal through its cognate receptor, Sucnr1 (Gpr91)^30^, we did not detect receptor expression in the ENS using our single-nucleus sequencing datasets. Instead, our genetic ablation studies demonstrated that succinate-mediated recovery requires tuft cells that express Sucnr1, suggesting that microbiota-derived succinate acts indirectly via epithelial and potential downstream immune pathways, resulting in ILC2 activation, leukotriene production, and acetylcholine release^30,31,50,51^. Consistent with this model, recent studies have demonstrated that succinate-producing bacteria activate tuft cell-dependent host-protective responses and confer resistance to *C. difficile* infection, further supporting a broader role for succinate in tissue repair and adaptation in response to microbial perturbation^52^.

While the ENS regenerates at a steady state from mature glial cells in zebrafish, for example^53^, in mammals, neurogenesis or neuronal regeneration does not occur in adults, except in the olfactory bulb and hippocampus^54,55^. The mechanisms of neuronal recovery described here could represent an evolutionary adaptation that sustains organismal fitness by regenerating neurons damaged by environmental perturbations such as toxins, infections, or inflammation^4,5^. The microbiota perturbation model studied here differs from paradigms involving overt tissue injury, such as benzalkonium chloride treatment, diphtheria toxin-mediated neuronal ablation, and severe colitis, which are associated with tissue damage and epithelial barrier disruption^56^. In contrast, Abx-induced dysbiosis results in neuronal and glial loss without overt tissue destruction, suggesting that the recovery responses identified in this study operate within a physiologically intact tissue environment. This distinction may explain why neuronal recovery following severe injury has generally been reported to be limited^13,15,42^ and suggests that microbiota-dependent recovery represents a specialized adaptation to reversible environmental challenges. Taken together, our findings identify microbial species and metabolites that promote recovery of the enteric nervous system and provide a mechanistic basis for how microbial signals stimulate neuronal regeneration in adulthood, with potential implications for post-infectious neuropathies and in regenerative medicine.

## Author contributions

B.A. conceived the study, designed, performed, and analyzed experiments, interpreted the data, and wrote the manuscript. I.M. analyzed the 16S rRNA sequencing, metabolomics, and initial snRNA-seq datasets and contributed to computational analyses and data interpretation. M.R.P. analyzed the snRNA-seq and scATAC-seq datasets and contributed to computational analyses and data interpretation. D.B. analyzed the time-series snRNA-seq dataset and contributed to computational analyses and data interpretation. J.C. performed and analyzed experiments and contributed to mouse treatments, ENS imaging and quantification, snRNA-seq experiments, and 16S rRNA sequencing. J.D. performed and analyzed experiments and contributed to ENS imaging and quantification, as well as intestinal functional measurements. H.L. performed and analyzed experiments and contributed to ENS imaging and quantification. S.Q. performed targeted SCFA metabolomics analyses and generated bacterial mutants with assistance from C.J.G. Y.A. contributed to mouse treatments, tissue dissections, and metabolomics sample preparation. D.Y.K. performed and analyzed experiments and contributed to imaging. S.A. assisted with experiments and imaging under the guidance of V.P. V.P.S. contributed to image analysis. L.D. and G.G. assisted with mouse treatments and tissue dissections under the guidance of I.M.C. M.A.S. assisted with snRNA-seq experiments. G.P.D. assisted with microbiological experiments. C.G. provided guidance and bacterial mutant strains. D.M. conceived, supervised the study, and co-wrote the manuscript. All authors read and approved the manuscript.

## Acknowledgements

We thank A. Rogoz and Luigi Bianco for maintenance of germ-free mice; A. Bilate for the germ-free rederivation of the glial reporter line; T. Ahrends, B. Reis, J.P. Truman for their help with flow cytometry and FACS. M. Fischbach (Stanford) for insightful suggestions on the metabolomics data; E. Gerrick and M. Horowitz (Stanford) for kindly providing the *T. musculis*; R. Hamnett and J. A. Kaltschmidt (Stanford) for their suggestions on structural analysis of the ganglia; B. Fait and N. Heintz (Rockefeller) for providing INTACT mice, Jacob von Moltke (WashU) for providing Pou2f3creERT2-ires-GFP mice, Ken Cadwell (Penn) for MM12+3 defined consortia mice, J. Faith (Mt. Sinai) for bacterial isolates of *B. vulgatus* and *B. pseudolongum*, M. Koenen and A.Kodra for their insights on nuclei sequencing protocol, and H. Alwaseem and M. Isay-Del Viscio for fecal metabolomics instrumentation and analysis at the Rockefeller University Proteomics Resource Center; Helen Duan and The Rockefeller University Sequencing Core for their help with sequencing 10X libraries and additional Rockefeller University employees for continuous assistance; and Zeny A. (*Felis catus*) for emotional support. Finally, we gratefully acknowledge the mice used in this study, which enabled these discoveries. Metabolomics data were generated by the Proteomics Resource Center at Rockefeller University (<u>RRID:SCR_017797</u>) using instrumentation funded by the Sohn Conferences Foundation and the Leona M. and Harry B. Helmsley Charitable Trust. Confocal microscopy was performed at the Rockefeller University’s Bio-Imaging Resource Center (RRID:SCR_017791). V.P.S. is supported by the Beckman Foundation award, AWD00000407. This work was supported by a grant from the Simons Foundation (718232, B.A.), H.H.M.I. Hanna H. Gray Fellowship (GT15986, B.A.), NIH R01DK126407 (D.M.), Food Allergy FARE/FASI Consortium (D.M. and I.M.C.). D.M. is an investigator of HHMI.

## Methods

### Mice

Mice were maintained at the Rockefeller University animal facilities under specific pathogen-free or germ-free conditions. In-house C57BL/6 or Jackson Labs C57BL/6J (000664) mice were used as wild-type mice, unless otherwise noted. Baf53cre (Tg(Actl6b-Cre)4092Jiwu/J, Jackson 027825); Plp1creERT (B6.Cg-Tg (Plp1-cre/ERT)3Pop/J, Jackson 005975); Sox10creERT2 (CBA;B6-Tg(Sox10-icre/ERT2)388Wdr/J, Jackson 027651); GFAPcreERT2 (B6.Cg-Tg(GFAP-cre/ERT2)505Fmv/J, Jackson 012849); Rosa26^DTA^ (B6.129P2 *Gt(ROSA)26Sor^tm1(DTA)Lky^*/J, Jackson 009669); Rosa26INTACT (B6.129-*Gt(ROSA)26Sor^tm5.1(CAG-Sun1/sfGFP)Nat^*/MmbeJ, Jackson 030952) transgenic mice were purchased and maintained under specific pathogen-free conditions on a standard chow diet in our facility. Pou2f3creERT2-ires-GFP (B6(129S4)-*Pou2f3^tm1.1(cre/ERT2)Imt^*/J., Jackson 037511) was obtained from Jacob von Moltke (WashU) and maintained at our standard SPF facility; Ser93 Sox10creERT2: Tg(Sox10-icre/ERT2)93Vpa, MGI ID: 5910373 (Vassilis Pachnis, The Crick) were housed at the Crick; GsdmE(-/-)(C57BL/6-*Gsdme^em1Hrlu^*/J, Jackson 040221) mice were housed at Harvard Medical School (Isaac M. Chiu) for the first set of experiments and were subsequently purchased directly from The Jackson Laboratory for independent replication studies. Genotyping was performed according to the protocols established for the respective strains by The Jackson Laboratory using the Transnetyx genotyping services. C57B6 and Swiss-Webster germ-free mice were purchased from Taconic Biosciences and maintained on a sterilized and autoclaved mouse diet (Cat:5021, LabDiet). The *Plp1^INTACT^* line was re-derived in a germ-free isolator and maintained on autoclaved food and water *ad libitum*. All mice were used starting at 7-12 weeks of age unless otherwise noted and housed under a 12-hour light/12-hour dark cycle with food and water *ad libitum*. Male or female sex-matched mice were pooled across all experiments because no significant differences between the sexes were observed in any of the parameters analyzed. Because of the limited number of mice interbred in our facility, not all experiments were performed with littermates. We did not control for the estrous cycle. All mice were handled in accordance with protocols approved by the Institutional Animal Care and Use Committee (IACUC) and the ethical guidelines of the National Institutes of Health (NIH) Guide for the Care and Use of Laboratory Animals.

### Antibiotics treatment

Five milligrams of ampicillin (Sigma-Aldrich) per mouse was administered via 100 µL oral gavage for 5-14 days, as indicated in the experiment. An Abx cocktail of 6.25 mg/ml of vancomycin Sigma-Aldrich), 6.25 mg/ml of ampicillin (Sigma-Aldrich), 12.5 mg/ml of metronidazole Sigma-Aldrich), and 12.5 mg/ml of neomycin (Sigma-Aldrich) (VAMN) was administered to *Salmonella ΔspiB*-treated mice for five days via oral gavage (100ul/mouse) before fecal microbiota transfer. Abx administration to GF mice was performed in sealed positive-pressure isocages with 1 g/L ampicillin in vacuum-sterilized (Corning) drinking water for seven days.

### Tamoxifen treatment

Tamoxifen (Sigma, T5648) was dissolved in sterile corn oil and ethanol (10%) to a concentration of 50 mg/ml. Plp1creERT mice received 7.5 mg/ml tamoxifen by oral gavage for induction. We observed increased off-target effects in neurons following multiple doses of Tamoxifen in the Plp1creERT line, thus limiting fate-mapping experiments to a single injection unless otherwise noted. Sox10creERT2 mice were gavaged with 7.5 mg/ml tamoxifen, and GFAPcreERT2 mice were gavaged with 12.5 mg/ml twice, as indicated in the experiment. Pou2f3creERT2 mice were gavaged with 5mg/ml tamoxifen. Ser93 Sox10creERT2 mice received 100 ug/g of body weight or 5 uL/g of body weight of a 20 mg/mL solution by oral gavage.

### Fecal microbiota transfer

Fresh feces (two pellets) from age-matched (or litter-matched whenever available) SPF mice were dissolved in 1 ml PBS and filtered with a 70 μM cell strainer. Oral gavage of 200 μl/mouse fecal mixture was performed, and mice were kept in cleaned cages for the experiment. *Salmonella ΔspiB-treated* mice received five days of VAMN Abx cocktail before fecal microbiota transfer as described above. For dysbiotic FMT transfers, fecal pellets collected on the last day of treatment with Abx or *Salmonella spiB* were frozen at −20°C, dissolved in PBS, and administered to the same mice, as described above.

### Salmonella *spiB* infection

Infections with *Salmonella ΔspiB* were performed as previously described^2^. Briefly, mice were pretreated with a single oral dose of streptomycin (20 mg/mouse in 100 ml of DPBS) 18-24 hours before infection. The mice were then orally inoculated with 10^9^ CFU of Salmonella *spiB*. A single aliquot of *Salmonella ΔspiB* strain was grown overnight in 3 mL of LB at 37°C with agitation. Bacteria were subcultured (1:300) into 3 mL of LB and incubated for 3.5 hours at 37°C with agitation. The pellet was resuspended to the final concentration in 1 mL of DPBS. Bacteria were administered by oral gavage at a total volume of 100 μL per mouse.

### Bacterial cultures and colonization of GF mice

Oligo-MM12 mice were maintained as gnotobiotic breeding colonies, with the defined microbial consortium transmitted vertically from dams to offspring. Mice aged 8-12 weeks were used for comparisons of enteric neuron density with age-matched GF mice. For monocolonizations, bacteria were cultured in Brain Heart Infusion (BHI; Becton Dickinson 211059) medium supplemented with 5 µg/ml hemin (Sigma 51280) and 1 µg/ml vitamin K (Sigma 95271). Mouse isolates were cultured either aerobically for *Escherichia coli* (Mt1B1) and *Enterococcus faecalis* (KB1)^20^ or in an anaerobic atmosphere of 85% nitrogen, 10% carbon dioxide, and 5% hydrogen for *Bacteroides vulgatus* (RU1) and *Bifidobacterium pseudolongum* (RU3). *Bacteroides sp.* 1_1_6 (*B. sp.* WT and Δ*mcm*)^28^strains were grown in the presence of 15 μg/ml thiamphenicol (Fisher Scientific). GF mice were colonized with *Escherichia coli* Mt1B1, *Bacteriodes vulgatus*, *Bifidobacterium pseudolongum*, *Bacteriodes sp.* 1_1_6 (*B. sp.* Δ*mcm*), and *Escherichia coli* Mt1B1::*mmdA* bacterial cultures grown to confluency by oral gavage of 200 μl/mouse in sealed positive-pressure isocages. Mice were sacrificed and harvested after seven–ten days of colonization for enteric nervous system staining and imaging, as indicated per the experiment. Colonization was assessed by qPCR with species-specific primers or by selectively plating on BBL Brain Heart Infusion (BHI) agar plates and Bacteroides Bile Esculin (BBE) agar plates.

### Tritrichomonas musculis colonization of GF mice

Frozen vials of *T. musculis* were thawed in a 37°C water bath and resuspended in 3 ml RPMI media without serum. The viability of protists was assessed using a hemocytometer under a bright-field microscope. The cells were centrifuged for 5 min at 100 × g and resuspended in PBS for mouse inoculation. GF mice were inoculated with *T. musculis* ∼100.000 cells per mouse by oral gavage in sealed isocages^26^.

### LPS treatment

Phenol-purified Lipopolysaccharides (LPS) from E. coli O127:B8 (Sigma-Aldrich) were dissolved in autoclaved drinking water and filter-sterilized. GF mice were gavaged daily for five days at a concentration of 5 mg/kg. The mice were harvested two days after treatment.

### 16S qPCR for assessment of bacterial colonization

Colonization of germ-free mice was verified by quantitative PCR (qPCR) using species-specific 16S rRNA gene primers. Fresh feces were collected for endpoint analysis. DNA extracted from fecal pellets using the Zymo Research Fecal DNA MiniPrep kit (D6010) and diluted 1:80 in nuclease-free water prior to analysis. qPCR reactions were performed using SYBR Green PCR master mix (Applied Biosystems, 43-687-02) on a QuantStudio 3 Real-Time PCR System (Applied Biosystems) with species-specific and universal bacterial 16S primers. Melt curve analysis was performed to confirm amplification specificity. Raw cycle threshold (Ct) values were used to assess the presence (colonization) of each bacterial strain across experimental groups. Universal 16S qPCR was performed in parallel to assess overall bacterial colonization

### Administration of bacterial metabolites

The following metabolites were administered to GF mice in sealed positive pressure isocages or previously SPF mice treated with Abx, as described above, in drinking water: 300 mM Sodium Chloride (Sigma-Aldrich), 150 mM Sodium Succinate (Alfa Aesar)^31^, 150 mM Sodium Propionate (Sigma-Aldrich), 150 mM Sodium Butyrate (Sigma-Aldrich), and 150 mM putrescine (Sigma-Aldrich). For GF mice, the dissolved metabolite solutions were filtered under a sterile hood using 500 mL vacuum filters (Corning).

### Construction of Bacteroides sp. 1_1_6 Δmcm mutant

The *Bacteroides sp. 1_1_6 Δmcm* mutant was generated using the allelic exchange strategy as described previously^28^. Briekly, approximately 1-kb regions klanking the *mcm* gene were amplikied and fused by fusion PCR using the primers 5’-CACAGGTTCATCTTCTCCCGCAGGACTTCACCGTATTCGG-3’ and 5’-CGATACCGTCGACTCGGAGGAACAGTAATCTGATCT-3’. The resulting fragment was assembled with the pExchange backbone by Gibson assembly, transformed into *E. coli S17*, and verikied by diagnostic PCR and sequencing. The verikied suicide vector was introduced into *Bacteroides sp. 1_1_6 Δtdk* by conjugation, followed by FUdR counterselection as previously described. Successful deletion was conkirmed by diagnostic PCR using the primers 5’-ACCGACTTCGGTATGGAAAAA-3’ and 5’-CCATTTCGGAAGAAAGCGG-3’.

### Heterologous expression of mmdA-scpA in *E. coli* Mt1B1

The coding sequences of *mmdA* (methylmalonyl-CoA mutase small subunit) and *scpA* (succinyl-CoA:propionate CoA-transferase) were amplified from genomic DNA of the donor strain *Bacteroides thetaiotaomicron* and cloned into the expression vector plasmid 82153-Pnifj-mmdA-scpA using Gibson assembly. The two genes were expressed as a synthetic operon under the control of the constitutive promoter Pnifj, with an independent ribosome binding site upstream of each coding sequence. An empty vector carrying the same plasmid backbone without the *mmdA-scpA* insert was used as a control. The resulting recombinant plasmid and the empty vector control were transformed into *Escherichia coli* Mt1B1 by electroporation. Electrocompetent cells were prepared by growing cultures in LB medium to an OD_600_ of 0.5–0.6, followed by three washes with ice-cold 10% glycerol. Electroporation was performed using 0.1-cm cuvettes at 1.8 kV. Cells were recovered in SOC medium for 1 h at 37°C and plated on LB agar supplemented with 25 μg mL^-1^ chloramphenicol and 100 μg mL^-1^ carbenicillin.

For metabolite production assays, *E. coli* Mt1B1 strains harboring either the *mmdA-scpA* expression plasmid or the empty vector were cultured in LB medium supplemented with 25 μg mL^-1^ chloramphenicol and 100 μg mL^-1^ carbenicillin. Overnight cultures were diluted 1:100 into fresh medium and grown for 18–24 h at 37°C. Expression and functional activity of the heterologous pathway were verified by targeted LC–MS quantification of succinate production, with empty-vector-transformed *E. coli* Mt1B1 serving as the negative control. For simplicity, the recombinant strain expressing the *mmdA-scpA* operon is referred to throughout the manuscript as *E. coli* Mt1B1::*mmdA*.

### EdU administration

0.8 mg/ml EdU (Thermo Fisher Scientific) dissolved in drinking water and given to mice ad libitum for 14 days. Fresh EdU water was supplied every two to three days.

### Intestinal transit time and length measurements

To measure the total intestinal transit time, mice were administered 6% carmine red (Sigma-Aldrich) by oral gavage, dissolved in 0.5% methylcellulose (prepared in sterile 0.9% NaCl). Transit time was recorded as the duration from oral gavage to the time at which the mice passed a fecal pellet containing carmine red. The experiments were concluded after 450 min, designated as the cut-off time for mice that had not passed fecal pellets.

To measure small intestinal transit time, mice received 200 uL of 25 mg/mL 70 kDa FITC-dextran (Sigma-Aldrich, 46945) in PBS via oral gavage and were sacrificed 12 minutes later. Upon sacrifice, the small intestine was dissected, carefully straightened, and the furthest point reached by FITC-dextran was identified with a fluorescent torch. Small intestinal transit was calculated as the distance traveled by FITC-dextran, normalized with the length of the intestine.

The mice were sacrificed to measure the small intestine length. The small intestine was cut distal to the stomach, carefully pulled out to detach it from the mesenteric tissue, and cut proximal to the ileocecal junction. The length of the whole small intestine was measured by laying it vertically alongside a measuring tape.

### Intestinal permeability assay

To assess intestinal barrier permeability, mice were deprived of food for 3 h before oral administration of a permeability tracer cocktail containing fluorescein isothiocyanate (FITC)-dextran (4 kDa; 20 mg per mouse; Sigma, FD4), and rhodamine B isothiocyanate (RITC)-dextran (70 kDa; 1.25 mg per mouse; Sigma, D1841). Three hours after gavage, mice were anesthetized with isoflurane, and blood was collected by retro-orbital bleeding. Blood samples were centrifuged at 1,500 × g for 10 min to isolate plasma, which was transferred to clear microcentrifuge tubes for analysis. Plasma concentrations of FITC-dextran and RITC-dextran were quantified using a fluorescence plate reader at excitation/emission wavelengths of 495/525 nm, and 555/585 nm, respectively.

### 16S sequencing

Fresh feces were collected for endpoint analysis. Sample preparation for 16S rRNA sequencing was performed by following the 16S Illumina Amplicon protocol from the Earth Microbiome Project^57^. MiSeq 2 × 150 with a 15% PhiX spike was used to sequence the libraries. Sequence processing and downstream analysis were performed in R. In summary, assembly into amplicon sequence variants (ASVs) and taxonomic classification using the Silva database were carried out with Dada2 (v1.2.6). Taxa observed fewer than 10 times in at least 20% of samples were excluded from the analysis. ASV/OTU quantification and related analyses were conducted using Phyloseq(v1.42)^58^ and ggplot2(v3.5.1).

### Extraction of cecal content for LS-MS

The feces were flash-frozen in liquid nitrogen. Frozen feces stored at −70°C were freeze-dried. Dried feces were weighed, and 30–40 mg were mixed with 800 μL of 80% methanol, 20% RNase, and DNase-free water. The samples were vortexed and sonicated for 10 min in a high-power sonicator. Samples were centrifuged at 13 000 rpm at 4°C for 15 min. The supernatant was collected and stored at −70°C until submission for LS-MS.

### Untargeted metabolomics of cecal contents

Mouse fecal extracts were analyzed by LC-MS/MS (QExactive Plus orbitrap mass spectrometer coupled to a Vanquish UPLC System, Thermo Fisher Scientific). External mass calibration was performed every three days using a standard calibration mixture. Polar Metabolites were separated by HILIC Chromatography (ZIC-pHILIC 150 x 2.1mm column, EMD Millipore) @150uL/min) using an analytical gradient decreasing from 90% B to 40% B in 30 minutes (A: 20 mM Ammonium Carbonate in Water adjusted to pH 9.3 and B: 100% Acetonitrile). The solvents were of LCMS grade quality (Optima, ThermoFisher). Each sample was analyzed in both positive and negative modes in separate injections. All MS/MS data were collected using Data-Dependent Acquisition (Resolution for Full-Scan:70,000, Resolution for Data-Dependent MS/MS Scans: 17500). Generated data were queried against MZCloud using Compound Discover 3.3 (Thermo Fisher). For matched molecules for which in-house standards were available, retention times were compared to the standards. The peak intensities of the MS/MS data were used to produce plots using Metaboanalyst^59^. The heatmap was plotted in R using the ggplot2 (v3.5.1) and pheatmap (v1.0.12) packages.

### Quantification of SCFAs with HPLC-MS

We constructed standard curves for acetate, propionate, and butyrate using the Area Under the Curve (AUC) of authentic chemical standards at different concentrations to quantify propionate and butyrate. Approximately 20 mg cecal samples were resuspended in 200 µL of 50% MeOH (in H_2_O) and vortexed for 10 min (beads (0.1×0.55 mm) were added to disperse the cecal material). Then, the mixture was centrifuged, and 10 µL of supernatant was mixed with 190 µL of short-chain fatty acid (SCFA) derivatization solution (1 mM 2,2’-dipyridyl disulfide, 1 mM triphenylphosphine, and 1 mM 2-hydrazinoquinoline, all dissolved in acetonitrile). The resulting mixture was vortexed, incubated at 60 °C for 1 h, and then centrifuged at 21000 × *g* for 20 min. The supernatant was analyzed using an Agilent 1290 LC system coupled to an Agilent 6530 quadrupole time-of-flight (QTOF) mass spectrometer with a 130 Å, 1.7 μm, 2.1 mm × 100 mm ACQUITY UPLC BEH C18 column (Waters). We used the following solvent system: A: H_2_O with 0.1% formic acid; B: Methanol with 0.1% formic acid. One microliter of each sample was injected, and the flow rate was 0.35 mL/min with a column temperature of 40 °C. The gradient for HPLC-MS analysis was: 0-6.0 min, 99.5%-70.0% A; 6.0-9.0 min, 70.0%-2.0% A; 9.0-9.4 min, 2.0% A; 9.4-9.6 min, 2.0%-99.5% A. Peaks were assigned by comparison with authentic standards, and relative analyte concentrations were quantified by comparing their peak areas with those of internal standards. The concentrations of propionate and butyrate were calculated using a standard curve and normalized to cecal weight.

For the detection of succinate, ∼ 20 mg cecal samples were resuspended in 200 µL of 50% MeOH (in H_2_O) and vortexed for 10 min (some beads (0.1×0.55 mm) were added to disperse the cecal material). Then, the mixture was centrifuged at 21000 × *g* for 20 min, and the supernatant was analyzed using an Agilent 1290 LC system coupled to an Agilent 6530 quadrupole time-of-flight (QTOF) mass spectrometer with a 1.7 μm, 2.1 × 100 mm ACQUITY UPLC® BEH C18 column (Agilent). The following solvent system was used: A: H_2_O with 0.2% formic acid; B: Methanol with 0.1% formic acid. One microliter of each sample was injected, and the flow rate was 0.35 mL/min at a column temperature of 40 °C. The gradient for HPLC-MS analysis was: 0-4.0 min 99.5% A-30.0% A, 4.0-4.5 min 30.0% A-2.0% A, 4.5-5.4 min 2.0% A-2.0% A, 5.4-5.6 min 2.0%A-99.5% A, with the instrument parameters settings as follows: collision and cooling gas, high-purity argon (Ar); nebulizing gas, high-purity nitrogen (N_2_, 1.5 L/min); gas temperature, 350 °C; collision energy, 40 kV; mode, negative mode; MS scan range, m/z 100-1600. The concentrations were calculated using a standard curve and normalized to cecal weight.

### Intestine dissections

Mice were sacrificed, the intestine was detached from the mesenteric tissue, and the 4-5 cm proximal to the ileocecal junction was taken as the ileum. For the dissection of the muscularis, the above segment of the whole tissue was laid flat, serosa facing up, and the muscularis layer was carefully peeled off using curved forceps, as described in more detail here^60^.

### Immunohistochemistry

Immunohistochemistry analysis of the myenteric plexus was performed as previously described^60^. Briekly, samples were kixed in 4% PFA overnight, washed in 1X PBS, and permeabilized in 0.5% Triton X-100/0.05% Tween-20/4 μg heparin (PTxwH). Samples were then blocked for two hours in blocking buffer (PTxwH with 5% BSA and 5% normal goat serum) at room temperature with gentle agitation. The samples were incubated with primary antibodies in blocking buffer for two days at 4°C with gentle shaking. After primary antibody incubation, the samples were washed four times in PTxwH and incubated with secondary antibodies for two hours at room temperature. The samples were washed four times in PTwxH and mounted using Fluoromount-G mounting medium (Thermo Fisher Scientikic). The following antibodies were used for immunohistochemistry experiments: Anna1 (1:10,000 (from human sera stock; Gift of Dr. Vanda A. Lennon), SOX10 (1:400 Ab227680), and NAV2 (1:500 Invitrogen PA5-52851), and nNos (1:1000, Ab1376). Fluorophore-conjugated secondary antibodies were either H&L or Fab (Thermo Fisher Scientific) at a consistent concentration of 1:400 in the following species and colors: goat anti-rabbit (AF488/568/647) and goat anti-human (AF488/568/647).

#### Immunofluorescence staining of tuft cells

Terminal ileal tissue was prepared as Swiss rolls, embedded in OCT compound, and cryosectioned at 20 μm thickness. Tissue sections were blocked in 5% donkey serum containing 0.1% Triton X-100 in PBS at room temperature before incubation with rabbit anti-DCAMKL1 (DCLK-1) antibody (1:500; Abcam, ab31704), anti-EpCAM (1:500; Thermo Scientific 25-9326-42); overnight at 4 °C. After washing, sections were incubated with the anti-Rabbit fluorescent secondary antibody (Thermo Fisher Scientific) diluted in blocking buffer for 2 h at room temperature in a humidified chamber. Fluorescence images were acquired using an Olympus FV4000 laser-scanning confocal microscope, and tuft-cells per villus were quantified.

### Fluorescence *in situ* hybridization

Fluorescence *in situ* hybridization on the ileum myenteric plexus was carried out using the Advanced Cell Diagnostics RNAscope® Fluorescent Multiplex Kit (Bio-Techne, 323100) according to the manufacturer’s instructions. The muscularis layer was peeled, fixed overnight in RNAase-free 4% PFA at 4 °C, and washed in RNAse-free PBS. Then it was dehydrated by serial ethanol treatments in 50% EtOH for 5 min, 70% EtOH for 5 min, 100% EtOH for 5 min, and 100% EtOH for 5 min. After dehydration, the tissue was treated with hydrogen peroxide for 10 min at RT, then with RNAscope® Protease III for 30 min at 40 °C. The tissue was then incubated with 4-Plex customized probes (RNAscope® Probe − Mm-Nav2, 313531-C1, RNAscope® Probe - Mm-Plp1, 428181-C3) at 40 °C overnight. Following hybridization, the tissue was washed twice with RNAscope® Wash buffer and then sequentially hybridized with pre-amplifier and amplifier DNA (Amp1-FL, Amp 2-FL, and Amp 3-FL) at 40 °C for 30 min, 30 min, and 15 min, respectively. After amplification, the tissue was hybridized with RNAscope® HRP solution for designated channels for 15 min, with RNAscope® TSA Vivid dyes for 30 min, followed by HRP blocker for 15 min. Following Fluorescence *in situ* hybridization, the tissue was immunostained with the Anna1 antibody as described above, counterstained with DAPI (1:10,000; Thermo Scientific, 62248), and mounted with Prolong Diamond Antifade Mountant (Thermo Scientific, P36970) according to the manufacturer’s instructions.

### Neuronal imaging and neuronal density quantification

Z-stack images of whole-mount samples were acquired on a Zeiss LSM 780 AxioObserver microscope with a Plan-Apochromat 25x/0.8 NA objective lens. A minimum of 8-15 images were randomly captured from a whole-mount muscularis sample. The Fiji cell counter tool was used to count the number of cells in each field^61^. Anna1 images were quantified using Cellpose v3.0.7^62^.We did not observe significant differences between manual and Cellpose counting of Anna1 images. Intraganglionic Sox10+ cells were counted and depicted on the graphs as glia. These counts were then multiplied by a factor of 3.125 (for a 25x objective) to calculate the number of neurons per square millimeter (mm²). The average from eight or more images was then calculated and plotted, with each point on the graph representing an individual mouse. The percentage of fate-mapped or NAV2+ neurons was calculated as divided by the total number of Anna1+ cells per image.

### Neurons per ganglia and nearest neighbor cell-to-cell distance quantification

Large-field, tiled, two-channel, Z-stack images were acquired on a Zeiss LSM 780 AxioObserver microscope with a Plan-Apochromat 25x/0.8 NA objective lens. Each image was 1024 × 1024 pixels in XY with a voxel size of 0.55 <u>µ</u>m x 0.55 µm x 1.00 µm. For image processing, maximum-intensity Z-projections were generated for each image. czi image in Fiji v2.14.0/1.54k using a custom script. The ganglion boundary was manually marked in each image using the following criteria. A myenteric ganglion was defined as a continuous group of Anna1+ cells separated by less than 15 µm. Only the complete ganglia were counted per field of view. Therefore, the following ganglia were excluded: 1. Ganglia truncated, 2. No clear separation (> 15 µm) was noted between the last Anna1+ cell and the edge of the field of view. Single Anna1+ cells separated by 15 µm on all sides were counted as extraganglionic. The quantifiable ganglia were averaged for at least 10 images per gut segment per animal. The Z-projection images were imported into QuPath v0.5.1, and all subsequent processing was performed in batch using custom Groovy scripts. Neuronal soma within each ganglion was identified in the Anna1 channel with Cellpose v3.0.7, and BIOP Cellpose extension v0.9.3, with the following parameters: model=cyto3 and diameter=15 pixels. Partial neurons that touched the ganglion boundary were removed. Delaunay clustering with a centroid-to-centroid distance threshold=45 μm was used to identify the Anna1+ nuclei cluster in each ganglion. The following measurements from the QuPath project were exported in a CSV file: image name, ganglion annotation name, Delaunay minimum distance, Delaunay number of neighbors, and cluster size. Raw CSV data were analyzed in Excel using pivot tables. Anna1+ cells with a cluster size < 3 were filtered out. The total number of Anna1+ cells in each ganglion was calculated and averaged for each mouse. For glia per ganglion quantification, nuclei within each ganglion were identified in the Sox10 channel with Cellpose v3.0.7 and BIOP Cellpose extension v0.9.3, with the following parameters: model=cyto3 and diameter=10 pixels. Partial nuclei touching the ganglion boundary were removed. Delaunay clustering with a centroid-to-centroid distance threshold=35 µm was used to identify the Sox10+ nuclei cluster in each ganglion. Sox10+ nuclei with a cluster size < 3 were filtered out. The total number of Sox10+ cells in each ganglion was calculated and averaged for each mouse. The nearest-neighbor cell-to-cell distance was averaged for each ganglion. All plots and statistical tests were performed using GraphPad Prism v10.2.1.

### Nuclei isolation from the muscularis layer

Mice were sacrificed. The small intestine was carefully detached from the mesenteric tissue, and 4-5 cm of the ileum was laid flat on a pre-chilled metal dissection block. The muscularis layer was carefully removed using curved forceps, placed in pre-chilled Eppendorf tubes, and frozen on dry ice. Frozen samples were stored at −80°C for up to a few weeks or processed immediately. To isolate the nuclei, frozen muscularis samples were taken from − 80°C and immediately placed into pre-chilled gentleMACS C-tubes (Miltenyi Biotech) and chopped with scissors for 20 seconds in 2 ml Nuclei Extraction Buffer (1M Tris-HCl, 5M NaCl, 1M MgCl_2_, 0.1% Tween with 0.04%BSA). The samples were placed on a gentleMACS Octo Dissociator (Miltenyi Biotech) and run on the built-in m_muscle program three times. The samples were immediately placed on ice for 10 min, occasionally swirling every 2-3 minutes. The samples were filtered directly on 30 µM MACS SmartStrainers (Miltenyi Biotech) and centrifuged at 500 × g for 5 min to isolate the nuclei. The supernatant was washed twice with nucleus-wash buffer (1% BSA in PBS). Additional iodixanol purification was used for the snRNA-seq experiment, as described in the section below.

### Nuclei Flow Cytometry

The nuclei of Plp1^INTACT^ mice were isolated as described above. The Click-iT Plus EdU Alexa Fluor 647 Flow Cytometry Assay Kit (Thermo Fisher Scientific, C10634) was used according to the manual to visualize EdU+ cells by flow cytometry. DAPI was used to stain the nuclei. The samples were acquired using a BD Biosciences FACSymphony A5 Cell Analyzer. Gating strategies for Plp1^INTACT^ mouse muscularis nuclei flow with EdU were as follows: DAPI+GFP+EdU+. Data was analyzed using FlowJo v10.10.

### scATAC-seq

Mice were sacrificed, and the five cm distal ileal muscularis layer was dissected, snap-frozen on dry ice, and stored at −80°C until processing. Nuclei were isolated as described above. Nuclei were resuspended in nuclei wash buffer (2% BSA in PBS) with DAPI (1:10,000). GFP-positive (*Plp1^INTACT^*) nuclei, enriched by fluorescence-activated cell sorting using a BD FACS Symphony S6 cell sorter equipped with an 85-μm nozzle and operated at 45 psi. Single-cell ATAC-seq libraries were generated using the Chromium Single Cell ATAC Reagent Kit (10x Genomics) according to the manufacturer’s instructions and sequenced on an Element Aviti platform.

### Single-nuclei RNA-seq (snRNA-seq)

#### Steady-state, Abx-treatment, FMT datasets (unenriched; *Fig. 3* And related)

The mice were sacrificed, 5 cm of distal ileal muscularis per mouse was carefully peeled off, and the tissue was frozen at −80°C. Four mice were pooled per condition into pre-chilled gentleMACS C tubes and nuclei were extracted as described above by adding Protector RNAse Inhibitors (Milipore-Sigma) at a concentration of 0.2U/ μL to Nuclei Extraction buffer and 1U/μL to Nuclei Wash buffer. The following density-gradient purification was done for the extracted nuclei: After washing once with 500 μL Nuclei Wash Buffer with RNase inhibitors, the pellets were resuspended in Sucrose buffer with RNAseq inhibitors (1M Tris-HCl, 5M NaCl, 1M MgCl_2,_ 320 nM sucrose, 1U/ul Protector RNase inhibitors) and gently mixed with 50% Iodixanol solution. The sample was underlaid with 30% iodixanol solution using an 18-gauge needle, and again with 35% iodixanol solution (35%, 30%, 25%, bottom to top). The samples were centrifuged at 10000g for 20 min at 4°C. The nuclei pellet formed between the 35% and 30% layers was carefully collected with a pipette. The pellet was resuspended in Nucleus Wash Buffer with RNAse inhibitors and centrifuged for 5 min at 500 g. The nuclei pellet was gently resuspended in Nuclei Wash Buffer with RNAse inhibitors and counted with an INCYTO C-chip hemocytometer (VWR) using a 1:1 ratio of 0.4% Trypan blue solution. The samples were immediately processed with a 10X Genomics Chromium Next GEM Single Cell 3’ kit v3.1, according to the manufacturer’s protocol. The libraries were sequenced on the Illumina NovaSeq platform.

#### Enrichment of Plp1 and HuC/D+ cells for snRNA-seq (*Fig 5*. and related)

Mice were sacrificed, and the 5 cm distal ileal muscularis layer was dissected, snap-frozen on dry ice, and stored at −80°C until processing. Nuclei were isolated as described above. For HuC/D-positive nuclei enrichment, nuclei were blocked for 10 min on ice in staining buffer (ST-SB; 2% bovine serum albumin (BSA), 0.02% Tween-20, 10 mM Tris-HCl, 146 mM NaCl, 1 mM CaCl₂, and 21 mM MgCl₂) and incubated with Alexa Fluor 647-conjugated anti-HuC/D antibody (Abcam, clone EPR19098, ab237235; 1:200) for 30 min on ice. Nuclei were then pelleted (800 × g, 8 min) and resuspended in wash buffer (2% BSA in PBS) containing DAPI (1:10,000). GFP-positive (*Plp1^INTACT^*) or HuC/D-positive nuclei were enriched by fluorescence-activated cell sorting using a BD FACS Symphony S6 cell sorter equipped with an 85-μm nozzle and operated at 45 psi. An unstained control lacking the HuC/D antibody was included to establish the HuC/D-positive gate. DAPI-positive GFP- or HuC/D-positive nuclei were collected for downstream single-nucleus RNA sequencing. Sorted nuclei were immediately processed using the Chromium Next GEM Single Cell 3′ Reagent Kit v3.1 (10x Genomics) according to the manufacturer’s instructions. Libraries were sequenced on an Element Aviti platform.

### snRNAseq processing

#### Steady-state, Abx-treatment, FMT datasets (unenriched; *Fig. 3* And related)

The snRNA-seq fastq files were processed using the ‘count’ function from Cell Ranger (10X Genomics, Cell Ranger, version 7.0.1) to align reads to the mm10 mouse genome, assign to cells/barcodes, and then map to genes. Samples were read into R for further processing and visualization with Seurat (version 5.0.3). Cell were filtered if they had a feature count <200 or >4000 leaving 52,933 total cells (unenriched data) or 15,144 (enriched data). Data were normalized with SCTransform, with regression to percent mitochondria followed by dimension reduction with PCA and UMAP with the first 20 PCs. Clusters were then determined using nearest-neighbors and leiden clustering (resolution – unenriched 0.5/ enriched 0.3). The unenriched samples were filtered to cell types of interest by removing clusters with particularly high expression of mesenchymal markers like Myh11 or Kit, while preserving clusters with high expression of Plp1, Sox10, Elavl4, Snap25, and Mki67, leaving 19,274 cells. Samples were integrated using Harmony (version 1.2.0)^63^, and then UMAP and clustering were rerun on this integrated object. Marker genes for each cluster and differential analysis were performed using findMarkers in Seurat. GSEA tests were performed with clusterProfiler (version 4.10.1).

#### GF and *B.sp. Δmcm* colonization datasets (Fig. 5 and related)

The snRNA-seq fastq files were processed using the ‘count’ function from Cell Ranger (10X Genomics, Cell Ranger, version 9.0.1) to align reads to the mm10 mouse genome, assign to cells/barcodes, and then map to genes. Counts from ambient RNA were removed, and empty droplets were filtered using CellBender (version 0.3.2)^64^. The cell by gene matrix was reads into R using Seurat (v5.0.3) and doublets were marked using Scrublet (version 0.2.3)^65,66^. The Seurat object was then filtered to exclude cells with fewer than 1% of their reads from mitochondrial genes, those identified as singlets by Scrublet, and those with fewer than 200 genes, resulting in 6218 cells. The data were then normalized and scaled using Seurat functions (NormalizeData, FindVariableFeatures, ScaleData), followed by linear dimension reduction (RunPCA) and non-linear dimension reduction using the UMAP method (RunUMAP) with the first 19 PCs. Low-resolution clusters were then determined using the Seurat functions FindNeighbors and FindClusters with a resolution of 0.05. Gene markers for each cluster were identified using the FindMarkers Seurat function, and based on the top markers, cluster 4 (muscle cell markers) and cluster 5 (Kit+ cells) were removed for further analysis. The remaining cells (4999 cells) were then re-normalized and scaled with the same Seurat functions indicated above, using the first 20 PCs for the RunUMAP and FindNeighbors functions. Clusters within the filtered cells were identified using a resolution of 0.1.

### snRNAseq downstream analysis

Each cell was then annotated with both functional neuronal subtypes and enteric neuron subtypes as previously defined from <u>the small intestine</u>^36–38,41^. The label transfer was performed with the FindTransferAnchors and TransferData functions from Seurat with an annotated Seurat object from the reference dataset. For the Fig. 5 dataset, very few cells were predicted to be in the enteric neuron class 11 (ENC11) cell type, so these cells were removed from the reference before the label transfer. Only cells that had a predicted max score of 0.5 or higher were assigned high-confidence annotations (17 cells were ambiguous). To understand differences in cell numbers of each sample within each cluster, a Fisher’s exact test was used. Expression patterns of gene sets across individual cells were calculated with the AUCell R Bioconductor package (v1.26.0)^67^. Single-cell potency scores and classifications were determined by the CytoTRACE2 R package (version 1.1.0)^68^. Trajectory analysis was performed using the Slingshot R package (version 2.10.0)^69^. For the Fig. 3. dataset, the dataset was first subset to the non-neuronal cells, and slingshot was run without specifying a starting cluster. For the Fig. 5 dataset, this was done separately with either Branch A (ENC8-12) or Branch B (ENC1-7) enteric neuron cells as defined in the Marklund p21 Morarach *et al.* reference. Cells annotated as glial cells were excluded from the trajectory analysis. For Branch A, ENC8 cells were used as the starting point and Branches 10 and 12 were defined as separate end points, while for Branch B, ENC1 was used as the starting group of cells. To generate plots, Seurat was used for violin and UMAP plots, the ggplot2 R package (version 4.0.0) ^70^ was used for the point, line, density, and bar plots, the ggalluvial R package (version 0.12.5)^71^ was used for the alluvial plots, and pheatmap R package (version 1.0.12) was used for heatmaps.

### Analysis of sc-ATACseq

scATAC samples were mapped, peaks called, and reads counted with cellranger-atac (version 2.1.0) using the mm10 mouse reference genome. Samples were read into R using Signac package (version 1.13.0) (Stuart, 2021). Cells were filtered if they were in the top 2% for reads in blacklist, nucleosome signal and peak number, or bottom 2% for percent of reads in peaks, TSS enrichment or number of peaks. They were also filtered if they were found to be doublets as assayed by scDblFinder (version 1.16.0)^72^ with either the DbltFinder or Amulet approaches, leaving 2726 total cells. TSS activity was determined using Signac and cell type annotation was transferred from reference datasets as done for snRNAseq datasets. Pseudobulk analysis was performed using DESeq2 (version 1.42.1) via the Libra wrapper (version 1.0.0). Coverage plots were generated with Signac’s CoveragePlot function, with labeling of consensus peaks, and peaks that significantly change in enrichment.

## Data availability

The raw data files for snRNA-seq, scATAC-seq, and 16S rRNA on SRA, and the processed single nuclei, 16S, and metabolomics data on GEO, are in the process of being uploaded.

## Biological material availability

Non-commercially available materials are available upon request.

## Statistical analysis

Experiments were performed without investigator blinding. All data points shown on the graphs represent biological replicates. The experimental groups included at least three biological replicates. Graphs and statistics were generated using GraphPad Prism v10.3.1. All data are presented as ±. An unpaired two-tailed Student’s t-test was used for experiments with two experimental groups. Samples with more than two experimental groups were analyzed using one-way ANOVA and Dunnett’s or Tukey’s multiple comparison test as indicated in the kigure legends. Multiple unpaired two-tailed Student’s *t*-tests with the Benjamini, Krieger, and Yekutieli false discovery rate correction were used for the panel in Extended Data Fig. 2f. Signikicance levels indicated in the graphs are as follows: * p < 0.05, ** p < 0.01, *** p < 0.001, **** p < 0.0001.

---

**Extended Data Fig. 1:**
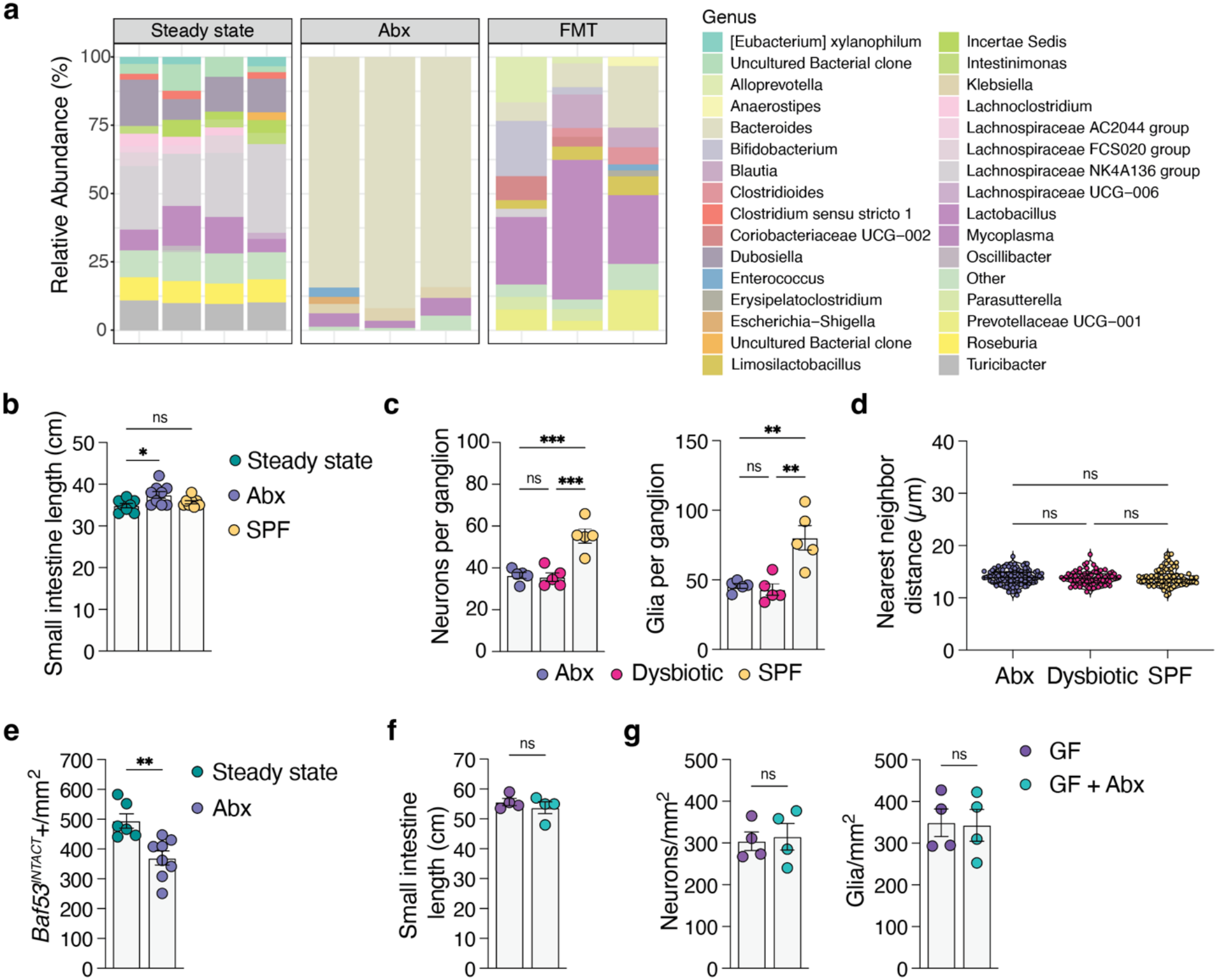
The effects of Abx and fecal microbiota transfer on ENS. **a**, The percent relative abundance of gut microbes at the genus level was analyzed by 16S rRNA sequences from fecal samples. *n =* 4,3,3. **b,** The length of the small intestine in mice was measured at the end of the experiment. Data are representative of at least two experiments. *n =* 8,9,8. **c,** Quantification of neurons per ganglion (left) and glia per ganglion (right) of mice treated with Abx or received dysbiotic or SPF FMT. *n =* 5,5,5. The same samples were analyzed, as shown in Fig. 1g. **d,** The distance between cells within the ganglia is represented by the nearest-neighbor distance. The same samples were analyzed, as shown in Fig. 1g. *n =* 30,30,30 cells in each group. **e,** Quantification of INTACT+ cells per mm^2^ in the Baf53cre^INTACT^ neuronal reporter line at steady state and five days after Abx. *n =* 6,8 **f, g,** GF Swiss Webster mice were treated with Anx in drinking water for seven days. Total changes in small intestinal length in **f** and quantification of neurons (left) and intraganglionic glia (middle) per mm^2^ area are shown in **g**. *n =* 4,4. The data are representative of at least two experiments except for panels **f** and **g**. All data are mean ±. One-way ANOVA with Dunnett’s multiple comparisons testing was used for **b,** One-way ANOVA with Tukey’s multiple comparisons testing for **c** and **d**. Unpaired two-tailed Student’s t-test was used for **e, f, g**.

**Extended Data Fig. 2:**
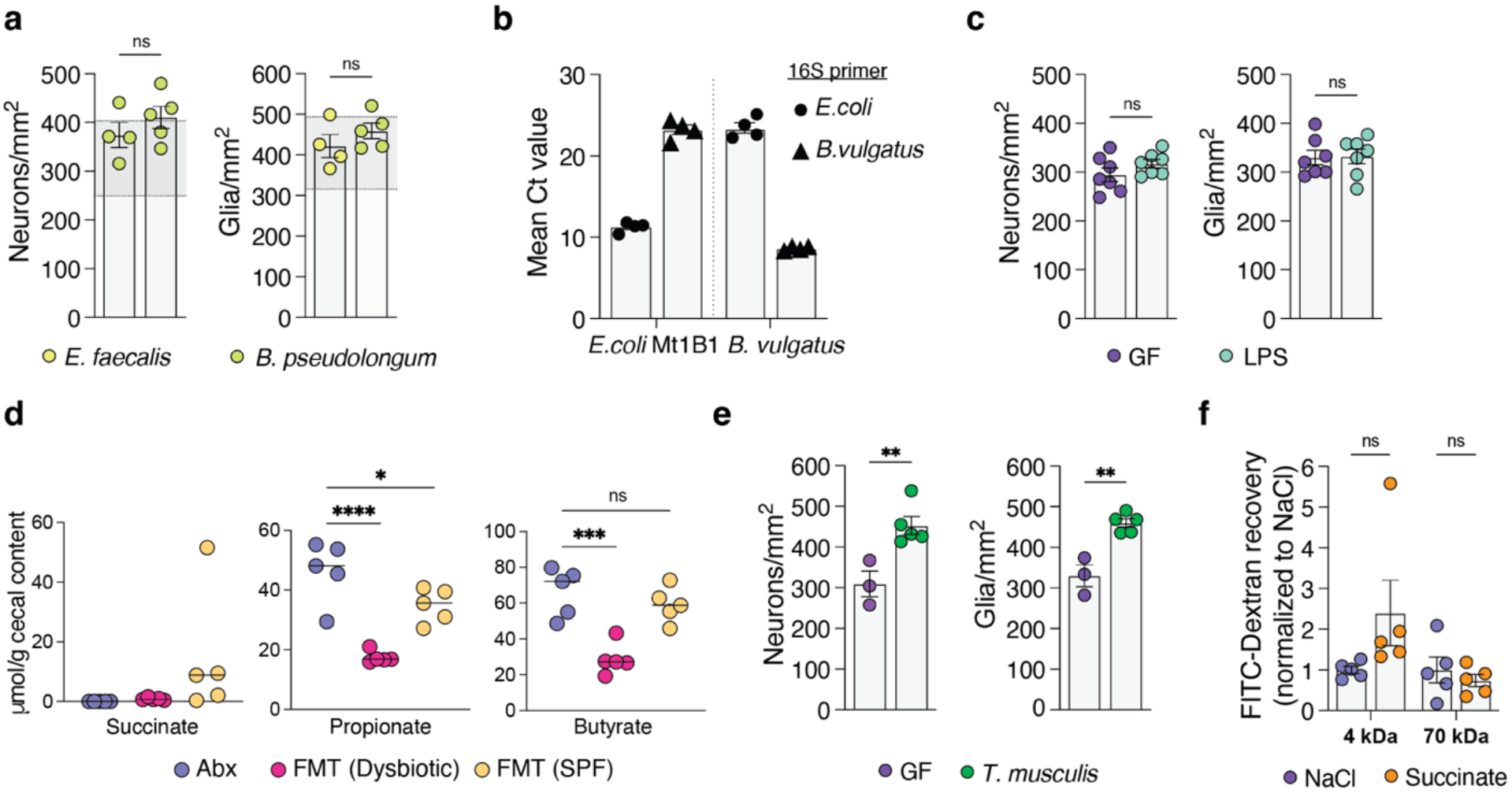
Characterization of the impact of succinate on ENS recovery. **a**, Quantification of neurons (left) and intraganglionic glia (right) per mm² of ileal myenteric plexus in GF Swiss Webster mice monocolonized with *E. faecalis* or *B. pseudolongum* for 7– 10 days. *n* = 4, 4 (*E. faecalis*) and *n* = 5, 5 (*B. pseudolongum*). Data are representative of at least two independent experiments. **b,** Quantitative PCR validation of monocolonization with *E. coli* and *B. vulgatus* using strain-specific 16S rRNA gene primers. Data are presented as mean Ct values. **c,** Quantification of neurons (left) and intraganglionic glia (right) per mm² of ileal myenteric plexus in GF Swiss Webster mice treated with oral gavage of *E. coli*-derived LPS for 5 days. *n* = 7. Mice were analyzed on day 7. **d,** Targeted LC–MS quantification of succinate and propionate levels in C57BL/6 mice treated with Abx or receiving fecal microbiota transplantation (FMT) from dysbiotic or SPF mice (*n* = 5 per group). Mice were analyzed 7 days after FMT. **e,** Quantification of neurons (left) and intraganglionic glia (right) per mm² of ileal myenteric plexus in GF C57BL/6 mice monocolonized with the protist *Tritrichomonas musculis* for 7 days. *n* = 3, 5. **f,** Assessment of intestinal barrier integrity by measuring relative serum levels of 4-kDa and 70-kDa FITC-dextran following oral gavage in Abx-treated mice receiving NaCl or succinate. Values were normalized to the mean NaCl group for each dextran tracer *n* = 5,5. All data are representative of two experiments except for panels **d, e, and f.** All data are mean ±s.e.m. Unpaired two-tailed Student’s t-tests were used for **a**, **c**, and **e.** One-way ANOVA, followed by Dunnett’s multiple comparisons test, was used for panel **d.** Multiple unpaired two-tailed Student’s *t*-tests with the Benjamini, Krieger, and Yekutieli false discovery rate correction were used for panel **f.**

**Extended Data Fig. 3:**
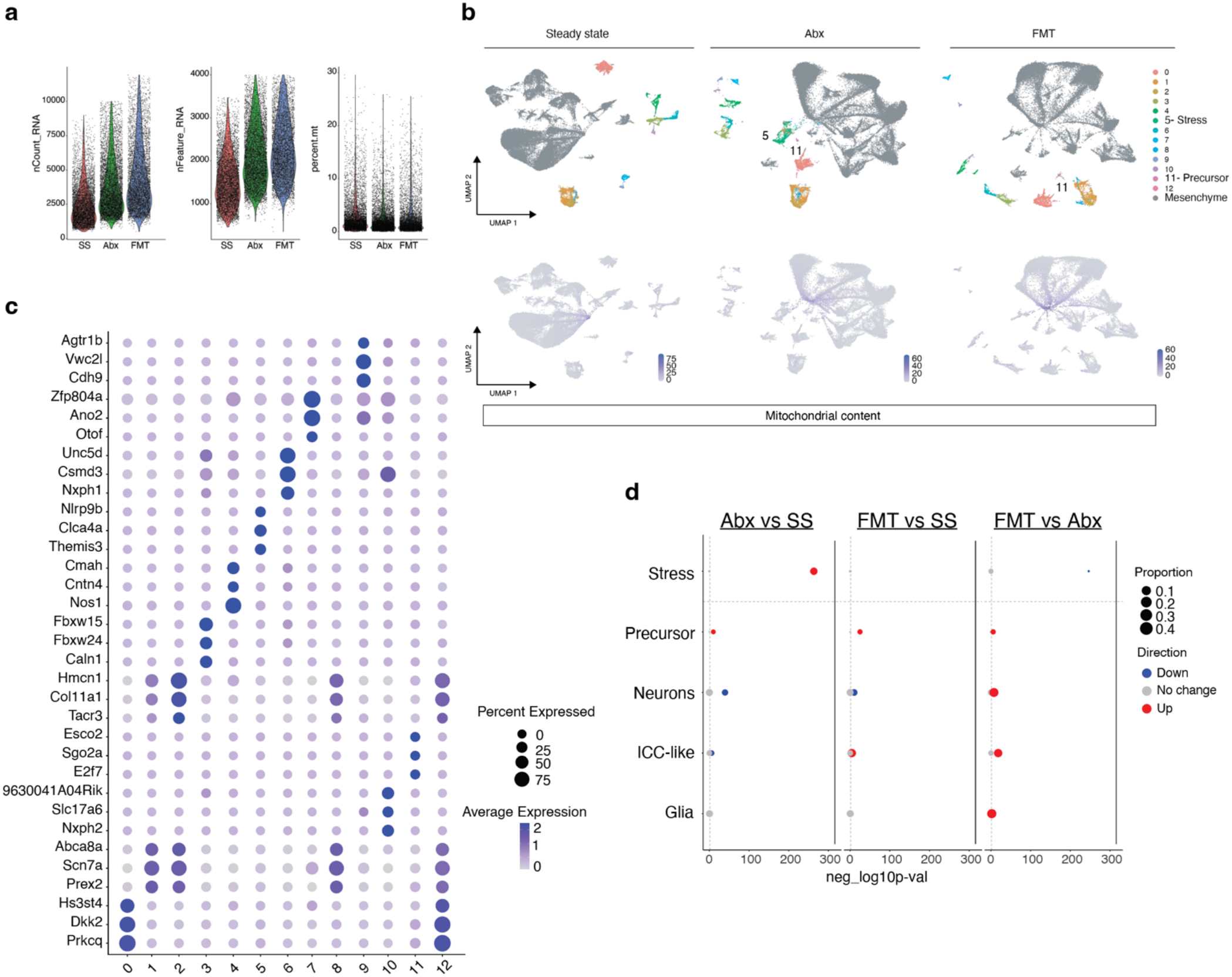
snRNA-seq characterization of the whole-tissue ileal muscularis layer. **a**, Quality-control (QC) metrics of the final snRNA-seq dataset following quality-control filtering, as described in the Methods. **b,** UMAP of whole-tissue ileal muscularis snRNA-seq datasets from steady-state (SS), Abx, and fecal microbiota transplantation (FMT) conditions. Upper panels show the subsetted populations analyzed in Fig. 3, and lower panels show mitochondrial gene signature scores for each library. **c,** Dot plot showing the top three differentially expressed genes for each cluster in the subsetted dataset shown in Fig. 3. **d,** Relative abundance of annotated cell clusters across SS, Abx, and FMT conditions, assessed by Fisher’s exact test.

**Extended Data Fig. 4:**
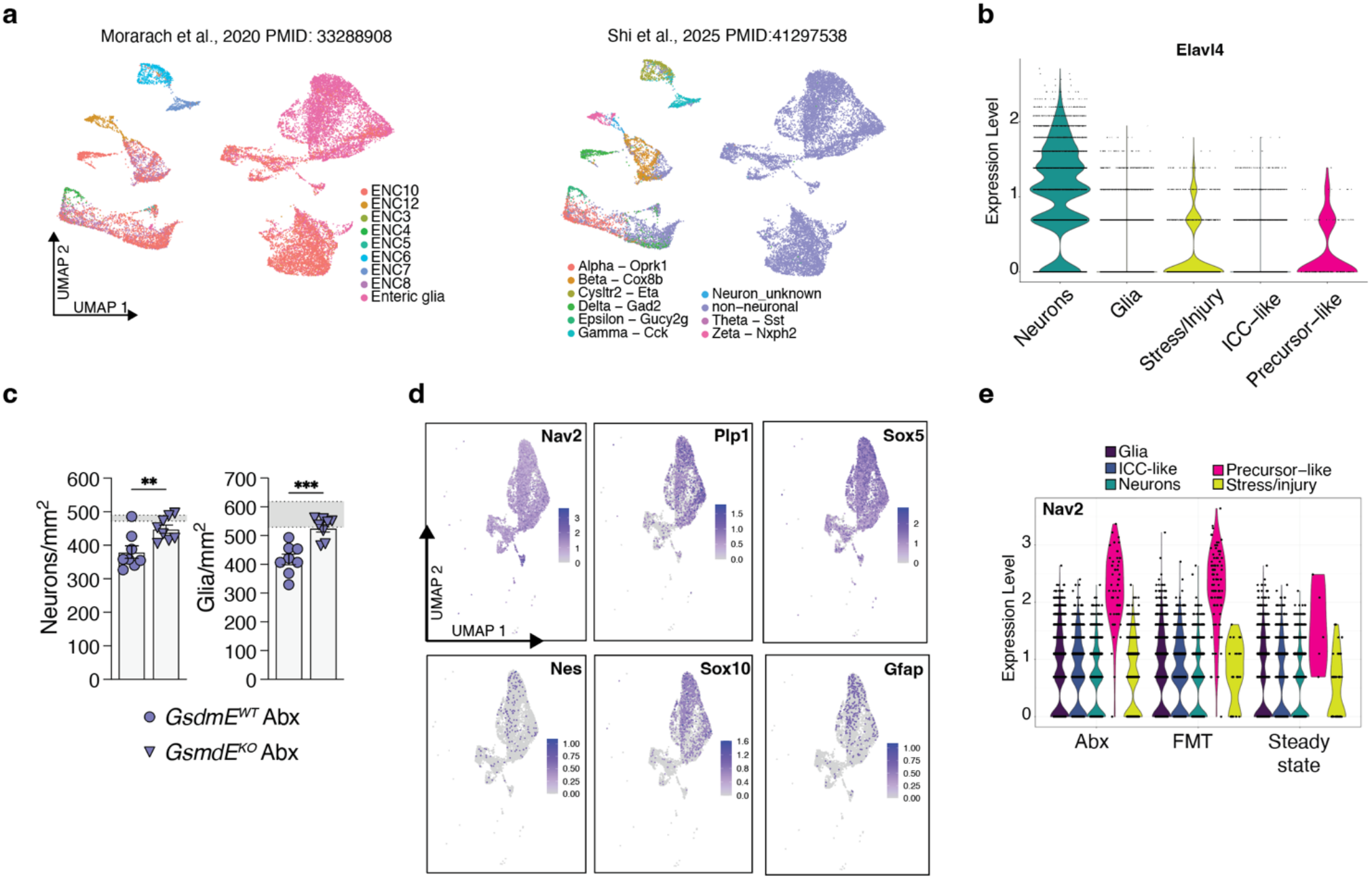
Additional characterization of injury- and recovery-associated cellular states. **a**, Label transfer from the published ENS scRNA-seq datasets of Morarach *et al.* (2020) (left) and Shi *et al.* (2025) (right). **b,** Violin plot showing *Elavl4* expression across the annotated cell clusters in the dataset shown in Fig. 3. **c,** Quantification of neurons (left) and intraganglionic glia (right) per mm² of ileal myenteric plexus in Gsdme+/+ (WT) and Gsdme−/− (KO) mice following 8 days of Abx treatment, matching the conditions used for the Abx snRNA-seq dataset. The gray-shaded region indicates the range of neuronal and glial densities in steady-state mice with the same genetic background. *n =* 8,8**. d,** Feature plots showing expression of representative glial marker genes across precursor-like, injury/stress-associated, and glial cell populations from the dataset shown in Fig. 3. **e,** Split violin plots showing *Nav2* expression across annotated cell clusters in the dataset shown in Fig. 3, separated by experimental condition.

**Extended Data Fig. 5:**
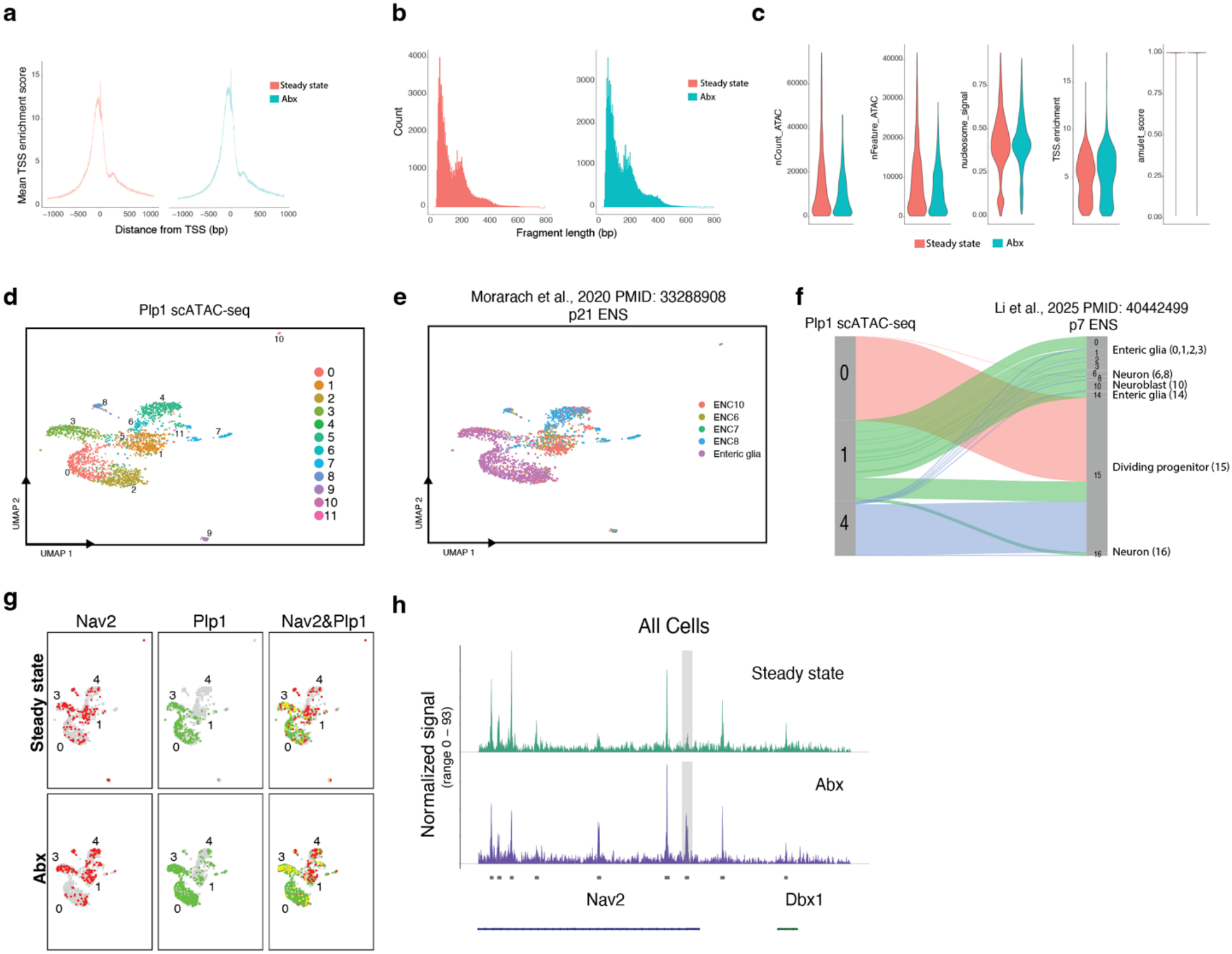
Characterization of the scATAC-seq dataset of Plp1+ cells at steady state and following Abx treatment. **a**, Mean transcription start site (TSS) enrichment scores for steady-state and Abx-treated scATAC-seq libraries. **b,** Distribution of fragment lengths across scATAC-seq libraries. **c,** Quality-control (QC) metrics of the scATAC-seq libraries following filtering. **d,** UMAP of the integrated Plp1+ scATAC-seq dataset from steady-state and Abx-treated mice. **e,** Cell-type annotation by label transfer using the p21 ENS scRNA-seq reference dataset from Morarach *et al.* (2020). **f,** Cell-type annotation by label transfer using the p7 ENS scRNA-seq reference dataset from Li *et al.* (2025). **g,** Gene activity scores for *Nav2* and *Plp1* across the integrated scATAC-seq dataset. **h,** Genome browser tracks showing normalized chromatin accessibility at the *Nav2* locus across all Plp1+ cells at steady state or Abx treatment. The gray shaded region indicates the differentially accessible peak shown in Fig. 4f.

**Extended Data Fig. 6:**
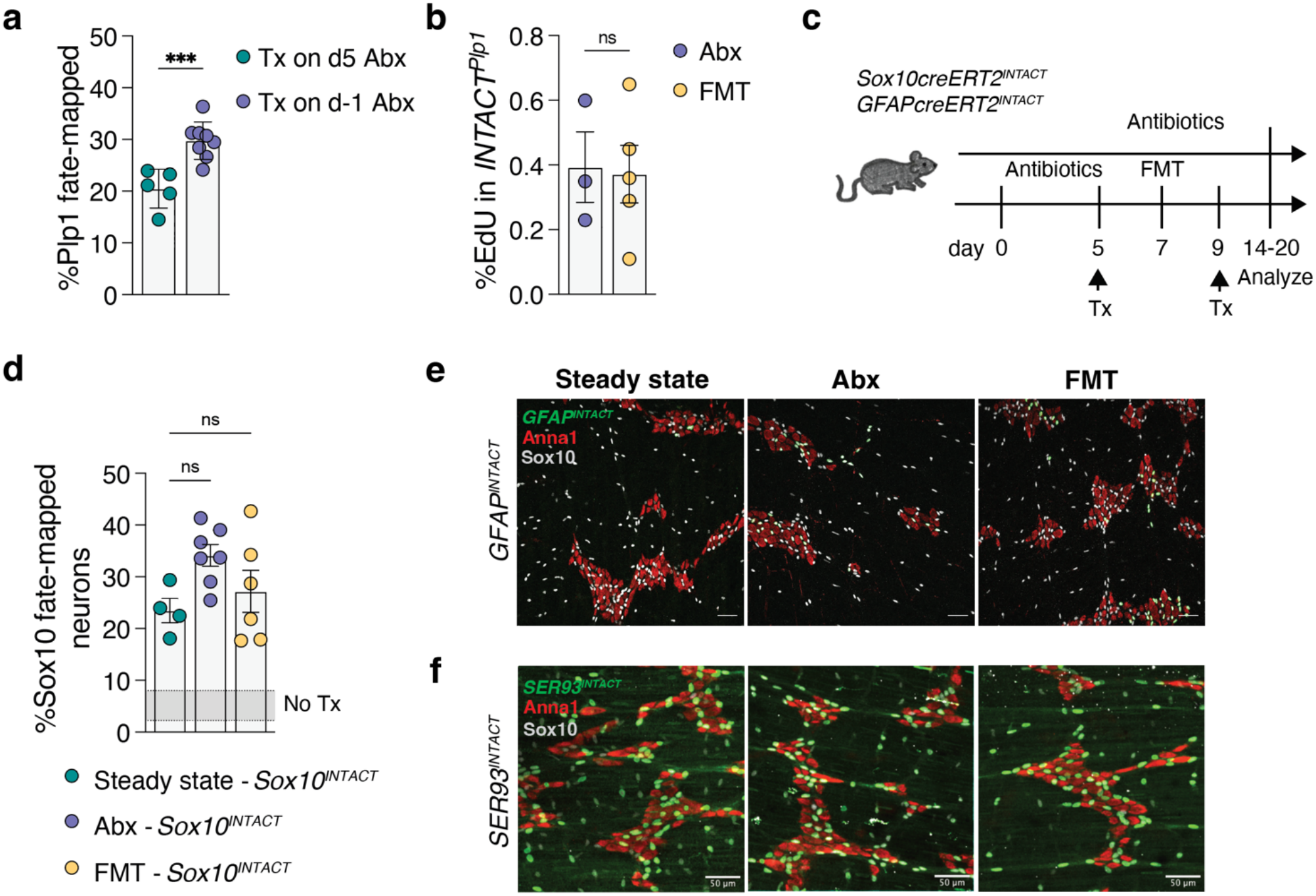
Glial fate-mapping strategies during ENS recovery. **a**, Quantification of the percentage of Plp1 lineage-traced neurons following tamoxifen administration either 1 day before (day −1) or on day 5 of Abx treatment. *n* = 5,8. **b,** Percentage of EdU+ *Plp1^INTACT^*+ cells in Abx and FMT groups, assessed by flow cytometry 3 days after FMT. *n* = 3, 4. **c,** Schematic of the experimental design for the inducible *Sox10^INTACT^* and *GFAP^INTACT^* glial fate-mapping experiments. **d,** Quantification of the percentage of *Sox10^INTACT^* Anna1+ cells among total Anna1+ neurons (left) and the density of *Sox10^INTACT^* +Anna1+ neurons per mm² of ileal myenteric plexus (right) 7–10 days following Abx or FMT *n* = 4,7,6. Steady-state mice received tamoxifen and were analyzed 24 h later. The gray shaded region indicates the distribution of baseline labeling in no-tamoxifen controls. Data are representative of at least two independent experiments. **e,** Representative confocal images of the ileal myenteric plexus from *GFAP^INTACT^* reporter mice immunostained for Anna1 and Sox10. No *GFAP^INTACT^* +Anna1+ cells were detected. Scale bars, 50 μm. Data are representative of at least two independent experiments. **f,** Representative confocal images of the ileal myenteric plexus from Ser93 *Sox10^INTACT^* reporter mice immunostained for Anna1 and Sox10. Scale bars, 50 μm. All data are presented as mean ± s.e.m. An unpaired two-tailed Student’s *t*-test was used for panels **a, b**. One-way ANOVA, followed by Dunnett’s multiple comparisons test, was used for panel **d**.

**Extended Data Fig. 7:**
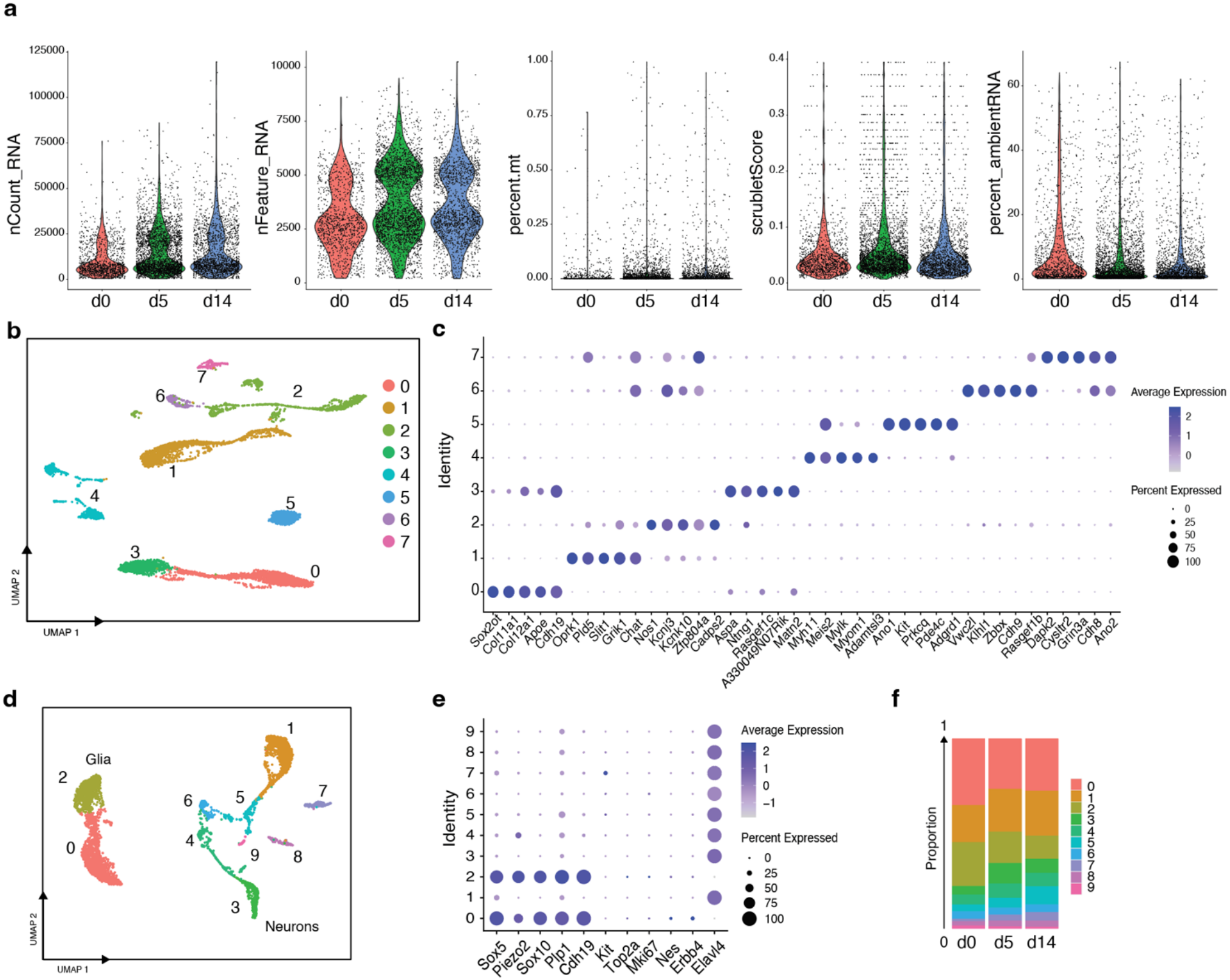
Characterization of the time-series snRNA-seq dataset from GF and *B.* sp*. Δmcm* monocolonized mice. **a**, Quality-control (QC) metrics for individual samples following quality-control filtering, including GF (day 0; germ-free), day five (d5), and day 14 (d14) monocolonization samples. **b,** UMAP of the integrated dataset showing the initial clustering prior to refinement. **c,** Dot plot showing the top five differentially expressed genes for each cluster in b. **d,** UMAP of the refined dataset used for downstream analyses following removal of clusters 4 and 5 from **b. e,** Dot plot showing the expression of representative marker genes associated with neuronal, glial, and ENS progenitor identities. **f,** Relative abundance of annotated cell clusters across experimental conditions.

**Extended Data Fig. 8:**
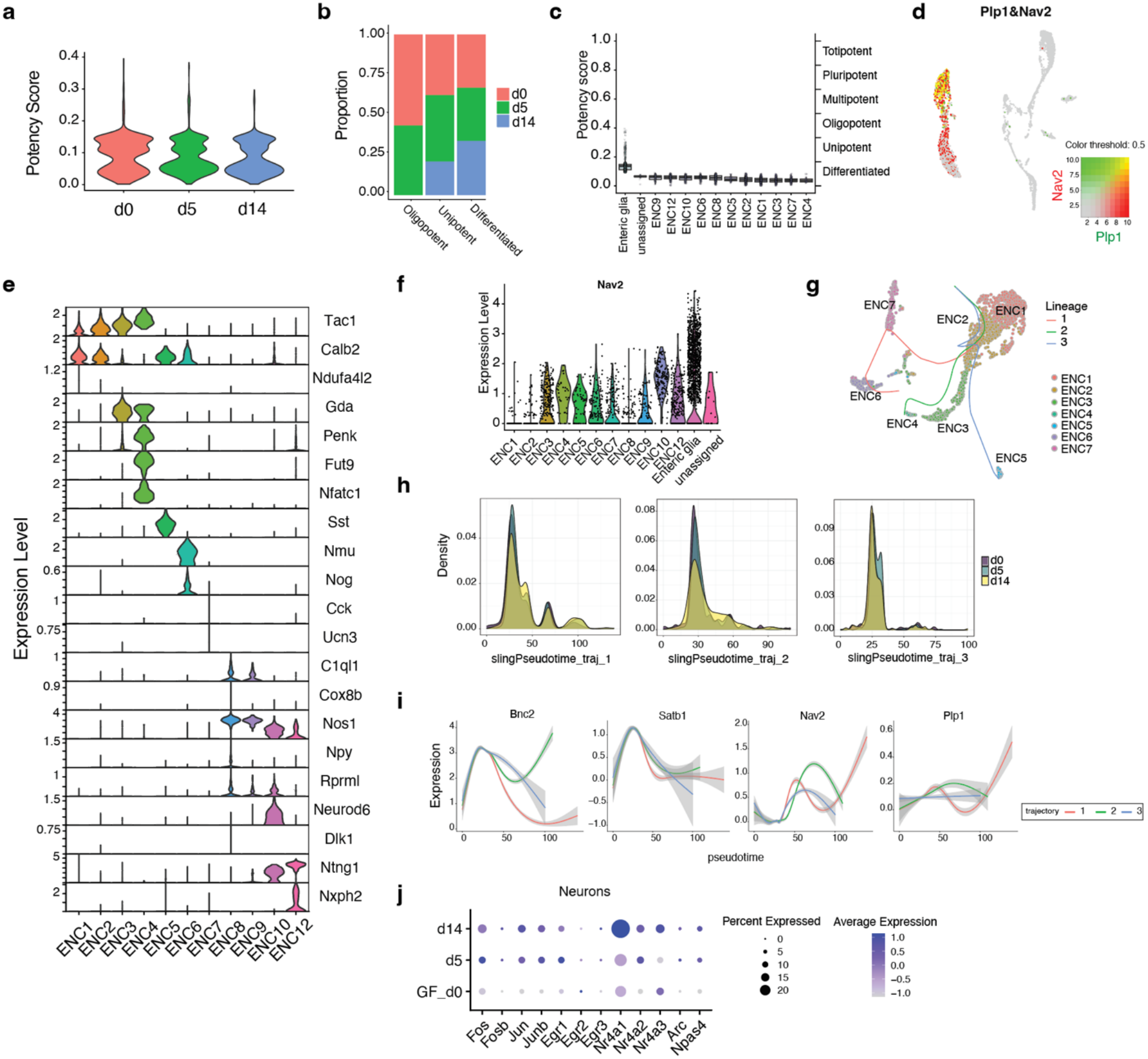
Additional characterization of developmental-like neuronal state transitions following *B. sp. Δmcm* monocolonization. **a**, CytoTrace2 developmental potency scores across individual experimental conditions. **b,** Proportion of cells classified as oligopotent, unipotent, or differentiated by *CytoTRACE* across experimental conditions, scaled to the total number of cells in each condition. **c,** Developmental potency scores across annotated cell clusters. d, Feature plot showing co-expression of *Plp1* and *Nav2* in the integrated dataset**. e,** Stacked violin plots showing the expression of representative marker genes defining ENS neuronal subtype clusters. **f,** Violin plot showing *Nav2* expression across annotated cell clusters. **g,** Slingshot trajectory inference of Branch B neuronal populations. **h,** Density plots showing the distribution of cells along the inferred Branch B trajectories shown in g. **i,** Expression of representative genes along the Branch B trajectories shown in **g. j,** Dot Plot showing the expression of immediate early genes (IEGs) across time points following monocolonization with succinate-accumulating *B. sp. Δmcm*.

**Extended Data Fig. 9:**
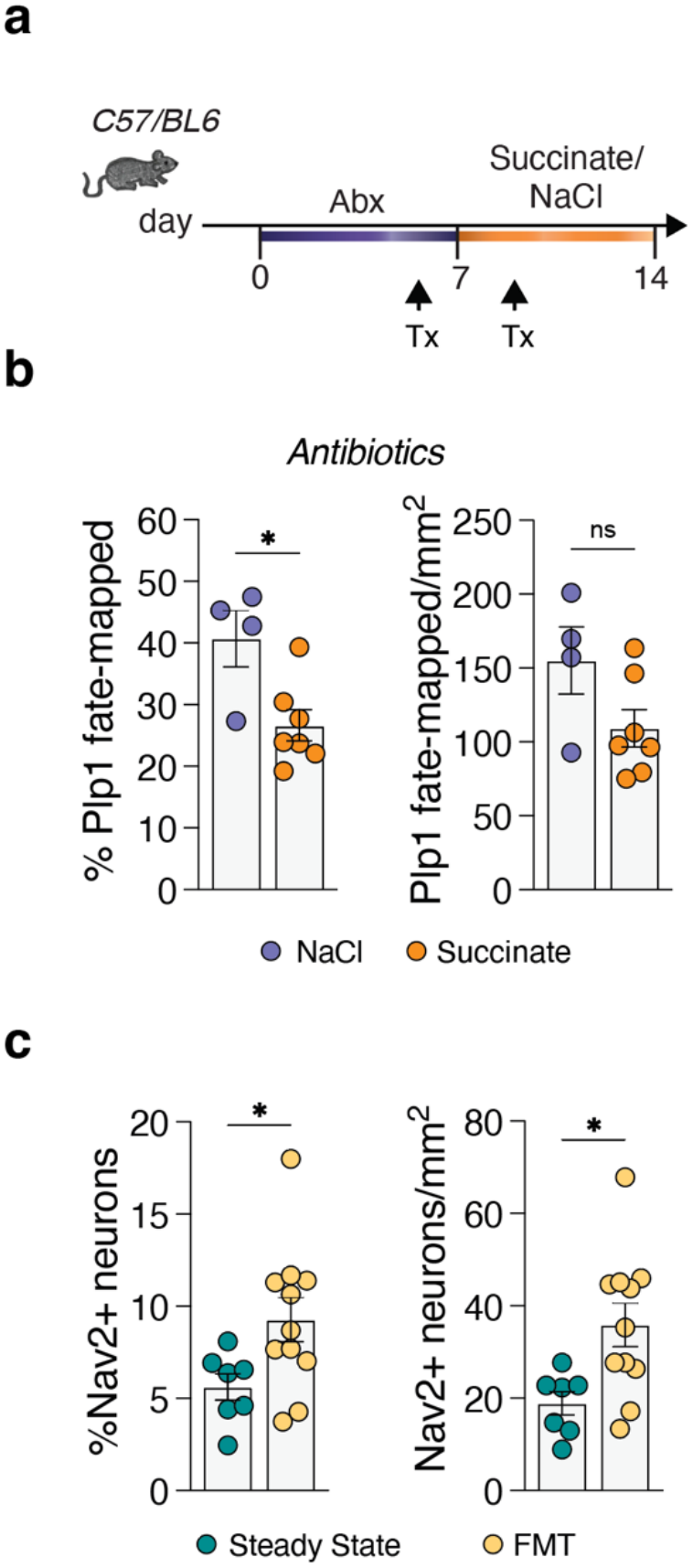
Plp1 fate mapping and Nav2-associated neuronal recovery following succinate treatment. **a**, Schematic of the experimental design for the inducible *Plp1^INTACT^* glial fate-mapping experiment during succinate-mediated recovery following Abx treatment. **b,** Quantification of the percentage of *Plp1^INTACT^* +Anna1+ fate-mapped neurons among total Anna1+ neurons (left) and the density of *Plp1^INTACT^*+Anna1+ neurons per mm² of ileal myenteric plexus (right) seven days following NaCl or succinate treatment. *n* = 4 (NaCl) and *n* = 6 (succinate). **c,** Quantification of the percentage of Nav2+Anna1+ neurons among total Anna1+ neurons (left) and the density of Nav2+Anna1+ neurons per mm² of ileal myenteric plexus (right) at steady state and following fecal microbiota transplantation (FMT). *n* = 7 (steady state) and *n* = 10 (FMT).

## Notes

### Competing Interest Statement

The authors have declared no competing interest.

### Summary of Updates

You are viewing Version 2, the most recent version of this article.

